# Depth-Dependent Contributions of Various Vascular Zones to Cerebral Autoregulation and Functional Hyperemia: An In-Silico Analysis

**DOI:** 10.1101/2024.10.07.616950

**Authors:** Hadi Esfandi, Mahshad Javidan, Rozalyn M. Anderson, Ramin Pashaie

**Affiliations:** Electrical Engineering and Computer Science Department, Florida Atlantic University, Boca Raton, FL, USA; Department of Medicine, University of Wisconsin-Madison, Madison, WI, USA; Geriatric Research, Education, and Clinical Center, William S. Middleton Memorial Veterans Hospital, Madison, WI, USA

## Abstract

Autoregulation and neurogliavascular coupling are key mechanisms that modulate myogenic tone (MT) in vessels to regulate cerebral blood flow (CBF) during resting state and periods of increased neural activity, respectively. To determine relative contributions of distinct vascular zones across different cortical depths in CBF regulation, we developed a simplified yet detailed and computationally efficient model of the mouse cerebrovasculature. The model integrates multiple simplifications and generalizations regarding vascular morphology, the hierarchical organization of mural cells, and potentiation/inhibition of MT in vessels. Our analysis showed that autoregulation is the result of the synergy between these factors, but achieving an optimal balance across all cortical depths and throughout the autoregulation range is a complex task. This complexity explains the non-uniformity observed experimentally in capillary blood flow at different cortical depths. In silico simulations of cerebral autoregulation support the idea that the cerebral vasculature does not maintain a plateau of blood flow throughout the autoregulatory range and consists of both flat and sloped phases. We learned that small-diameter vessels with large contractility, such as penetrating arterioles and precapillary arterioles, have major control over intravascular pressure at the entry points of capillaries and play a significant role in CBF regulation. However, temporal alterations in capillary diameter contribute moderately to cerebral autoregulation and minimally to functional hyperemia. In addition, hemodynamic analysis shows that while hemodynamics within capillaries remain relatively stable across all cortical depths throughout the entire autoregulation range, significant variability in hemodynamics can be observed within the first few branch orders of precapillary arterioles or transitional zone vessels. The computationally efficient cerebrovasculature model, proposed in this study, provides a novel framework for analyzing dynamics of the CBF regulation where hemodynamic and vasodynamic interactions are the foundation on which more sophisticated models can be developed.

**Author summary:** Blood vessels dynamically adapt to the mechanical forces exerted by circulating blood. Appropriate adaptive responses to changes in mechanical force are central to the optimal functioning of the cerebral blood flow (CBF) regulatory system, and include processes such as cerebral autoregulation, vasomotion, and neurogliovascular coupling. This adaptation is driven by intercellular interactions, primarily modulated by factors such as vessel wall tension, shear stress, and strain. As our understanding of the biophysicochemical principles of CBF regulatory system has advanced, computational studies have become more detailed and sophisticated, providing practical in-silico environments to investigate its dynamics and gain insight into the underlying biology. In this study, I propose a method to create a computationally efficient platform where the interactions of hemodynamics with vessel segments can be modeled and studied in an in-silico setting. This method can lay the groundwork for more sophisticated computational studies of the CBF regulatory system, where hemodynamics are core elements of the system operation and the model can represent a more realistic version of this system.

## Introduction

Autoregulation serves as the primary mechanism regulating CBF. This regulation is driven by the myogenic response in vessels[1]. As mean arteriolar pressure increases, the potentiation of the myogenic response works to maintain relatively constant blood flow throughout the vascular network. Additionally, during periods of increased brain activity, the inhibition of the myogenic response through neurogliavascular coupling (NGVC) leads to vasodilation and enhancement of blood delivery to activated regions— a process termed functional hyperemia (FH). Understanding the mechanisms involved in the autoregulation and NGVC and the tight interplay between them in health and pathological condition has attracted significant attention in recent year[1, 2]. Analyzing variables such as RBC flux and velocity, blood flow velocity, vessel diameter, and RBC line scan density has become a common practice in evaluating cerebral autoregulation[3, 4, 5, 6, 7, 8, 9, 10, 11, 12]. These physiological variables can be assessed quantitatively under static conditions including mild and stable anesthesia or in awake mice without potent and transient neuronal activity. Therefore, it is scientifically valuable to understand how these variables are influenced by factors such as cortical depth, vascular zone, disrupted myogenic response, and particularly mean arterial pressure level which varies across diverse experimental settings, especially due to variations of the heart rate[13].

Moving beyond the vascular-centric process of autoregulation[1], there is growing interest in investigating hemodynamics and vasodynamics across various vascular zones during FH to evaluate NGVC under pathological conditions[14, 15, 16, 17]. The diversity of vasodilatory mechanisms across various vascular zones is well-recognized [2, 18]. This diversity necessitates identifying the vascular zones where impaired NGVC could most detrimentally affect blood delivery. Controversies exist in in-silico studies aiming to quantify in which vascular zones NGVC-induced vasodilations contribute most significantly to blood delivery. For example, Gould et al. suggest that capillaries adjacent to feeding arterioles, now recognized also as transitional zone (TZ) vessels, are key contributors to hydraulic resistance[19]. Another study by Hall et al. calculated that a 6.7% dilation in cortical capillaries could account for 84% of the total blood flow increase seen during FH[20]. However, it is worth noting that in their study, vessels categorized as capillaries included the first four orders of bifurcated vessels from penetrating arterioles (PAs), which are now more specifically referred to as TZ vessels or precapillary arterioles. In contrast to these studies, Rungta et al.’s experimental and computational analysis of FH dynamics presented a different viewpoint proposing that vasodilation in higher-order capillaries significantly contributes to FH[21]. These differing interpretations underscore the challenges in quantifying these dynamics, particularly without considering the effect of factors such as the basal degree of constriction in vessels, cortical depth, intensity and period of neuronal activity, and mean arteriolar pressure.

Computational studies of autoregulation have predominantly concentrated on the dynamics of CBF regulation through time-domain analyses upon the application of vasoactive stimuli. These models often employ lumped mathematical models rooted in the principles of static autoregulation[22, 23, 24, 25, 26, 27, 28, 29]. Despite these efforts, a gap remained in the computational analysis of core aspects of static autoregulation, particularly within the cerebral vasculature. Static autoregulation is a vascular-centric regulatory process which functions by precisely transducing mechanical forces exerted on internal surfaces of blood vessels into mechanical forces exerted by mural cells on the outer surface of the lumen. This mechanism finely adjusts luminal diameter, ensuring uniform and consistent blood flow across all cortical depths even amidst substantial variations in intravascular pressure (IP) within the brain’s main feeding arteries. Therefore, both outputs (like vessel diameter and CBF adjustments) and inputs (mechanical forces exerted by the bloodstream on the vessel walls) are coupled hemodynamic and vasodynamic variables, integrated into this system. The tight coupling of vasodynamics and hemodynamics highlights the less explored potential of incorporating analytical approach to simulate these variables within cerebrovascular networks for studies focused on cerebral autoregulation [30]. To date, hemodynamics-vasodynamics analysis in cerebrovascular models have been constrained by the application of fixed boundary conditions and constant vessel diameters derived from segmented two-photon microscopy images[19, 31, 32, 33, 34, 35, 36, 37].

The main objective of this work is to examine how the potentiation and inhibition of the myogenic response across different vascular zones contribute to cerebral autoregulation and enhance blood delivery during FH. To accomplish this we adopted several justifiable simplifications and generalizations to provide an abstract yet insightful perspective on the key aspects of cerebral autoregulation using a simulated vasculature. We began by designing our in-silico cerebrovascular model and progressively refined it by incorporating essential morphological and mechanobiological characteristics of the mouse cerebrovasculature. The analysis focused on achieving optimal IP distribution across the network’s nodes and regulating vessel segment diameters, and therefore the resistance, to maintain relatively consistent blood flow in capillaries across all cortical depths, despite significant IP variations in main feeding arteries. This approach enabled us to computationally investigate the system’s input-output relationship across diverse cerebrovascular zones and analyze the necessary precision in these translations for ideal blood distribution.

## Results

### A: Design and evaluation of a cerebrovascular model

Cerebral blood flow is directly proportional to the IP gradient between the main cerebral arteries and veins, and inversely proportional to the blood flow resistance between them. This resistance is determined by factors such as the vessel diameter and rheological properties of the blood. To build the vascular model for the analysis of cerebral blood flow autoregulation and vasodilation during FH, we need reasonable estimations of basal internal vessel diameters pre-FH. For a vessel to form a suitable pathway for RBC movements, a force such as the intravascular pressure should be applied to its elastic structure to induce passive distention and create a tubular shape with a measurable diameter. In addition to the intravascular pressure, the force exerted by mural cells, muscular cells encasing the vessel’s lumen, including smooth muscle cells (SMCs) and pericytes, can cause active constriction which is another factor in determining vessel basal diameter. Thus, diameter of a vessel segment is the outcome of the dynamic equilibrium between these two opposing forces, the passive distension and active constriction. The magnitude of passive distension depends on the vessel’s distensibility and is modulated by circumferential stress applied to the internal vessel wall. Meanwhile, the extent of active constriction in each vessel segment depends on the concentration of contractile elements and is modulated by hemodynamic variables within the segment, including internal pressure and blood flow. Therefore, for accurate modeling of an elastic vessel segment that dynamically adjusts its diameter in response to internal hemodynamic variations, we need information about the structural composition of vessel walls and the way the muscular force is modulated by the hemodynamics. With this information in hand, we can calculate the equilibrium between these forces in each vessel segment based on internal hemodynamic values and use that to estimate the diameter of the segment and its resistance to blood flow.

It is technically challenging to obtain the information needed to model each vessel segment for two reasons: 1. The structural composition of vessel walls varies across different vascular zones and locations. To address this, we incorporated the morphological characteristics of mouse cerebral vasculature to design the initial framework of our in-silico vasculature, 2. Identifying the precise transfer function, the function that maps vessel’s internal hemodynamics to the muscular force imposed on the vessel, is currently infeasible due to several unknown principles that govern the calibration of this transfer function. There are multiple hypotheses on how vessels adapt their lumen diameters to hemodynamics. A widely accepted perspective suggests that each vascular segment independently seeks to balance the mechanical forces it faces (such as wall tension and wall shear stress), aiming for durability and minimizing energy expenditure in the process of blood flow regulation [38, 39, 40]. However, the specifics of these multivariable optimizations have remained unclear, including whether vascular cells autonomously manage this minimization process and establish the transfer function, or if intermediaries like astrocytes continuously aid in adjusting blood flow, participating in cerebrovascular adaptation to precisely meet metabolic demands and calibrate these transfer functions[41]. Despite these uncertainties, it is reasonable to assume that these processes collectively strive to autoregulate the blood flow in a healthy vasculature. This cause-and-effect assumption guided our approach in estimating the calibrated transfer function for every segment within our in-silico vasculature.

Therefore, in the subsequent parts of this section, we employ numerical and analytical methods to refine the cerebrovascular model and estimate the transfer function of all segments to enable functional autoregulation. The objective is to develop a computationally effective coarsely segmented cerebral vasculature with a calibrated transfer function for each segment that dynamically relates its internal hemodynamics to the vessel diameter. Within this network, the interplay between changes in the hemodynamics and vessel diameter, particularly when considering the existence of a delay in muscular force adjustment in response to hemodynamic variations, leads to a scenario where vessels cannot maintain a dynamic balance between passive and active forces. In a system with a negative feedback, any increase in the output triggers a response that decreases that output. In the CBF regulatory system, a rise in IP triggers active constriction, effectively reducing the IP as a form of a negative feedback. However, unlike the instantaneous passive distension, active constriction involves a sequence of events—from detecting changes in IP to inter-cellular interactions in mural cells that regulate myosin light chain phosphorylation—thereby delaying the adjustment of the muscular force. This strategy of integrating a delayed negative feedback, as seen in various biological regulatory systems for inducing autonomous oscillations[42], is similarly employed in the CBF regulatory system which generates oscillatory dynamics. Consider a PA experiencing an increase in IP. Although the negative feedback should counter this increase, the interplay between the delayed active constriction and instantaneous passive distension instead leads to an augmented diameter and, subsequently, elevated pressure. This process creates a temporary positive feedback scenario until the delayed adjustments in muscular force overcome the passive distension, reverting to its negative feedback role to curb the pressure increase and halt the dilation cycle. Once the constriction cycle initiates, the IP within the PA begins to drop immediately, yet the force applied by mural cells remains adjusted to peak IP levels. This condition prompts mural cells to progressively reduce the diameter until the delayed adjustment in muscular force is inadequate against passive distension, ending the constriction cycle and triggering the new oscillations of the vessel diameter—a phenomenon known as vasomotion. For the purposes of our analysis in Section B, we simplify the model by omitting this delay. This simplification allows us determine the steady-state diameter of vessels.

Currently, realistic cerebrovascular models, which are derived from segmented images generated by two-photon microscopes, are commonly used to analyze cerebrovascular hemodynamics. Simulating a high-resolution segmented cerebrovascular model requires significant computational power, especially when used for temporal and multi-scale quantitative analysis[43]. Fig 1(a) shows an example of a coarsely segmented vasculature model suitable for multi-scale studies and for the modeling of each vascular segment. While these models are useful in some studies, we encountered several challenges that made them impractical for our specific research needs. First, the inherent spatial limitations of segmented vasculature typically result in a large number of boundary nodes throughout these networks. This situation requires estimating the unknown boundary conditions for less critical nodes, using physiologically informed conditions applied to principal nodes, to address the ill-posed nature of hemodynamic analysis within these networks[31]. Such a process potentially diminishes the precision of hemodynamic analysis in cerebrovascular model, which is the core aspect of our analysis. Moreover, in working with cerebrovascular models, precise identification of vascular zones and types of mural cells that regulate each segment’s diameter is essential. However, reaching this detailed level is difficult in realistic coarsely segmented cerebrovascular models without specific information about the types of mural cells encasing each vessel segment. This difficulty arises from the inability to distinguish between PAs and TZ vessels, as well as between TZ vessels and capillaries, especially deeper in the cortex, when relying solely on vessel diameter and vascular organization. Additionally, our preliminary hemodynamic analyses conducted on realistic coarsely segmented vasculature models, resulted in significant inhomogeneity in hemodynamic variables across the network, deviating from reality. This discrepancy underscored the need to develop a simplified yet computationally effective model. To address the complexities inherent in coarsely segmented realistic microvasculature, we developed an artificial vasculature depicted in Fig 1(b). Design principles of this model are discussed in detail in Sec A2.

**Fig 1.**
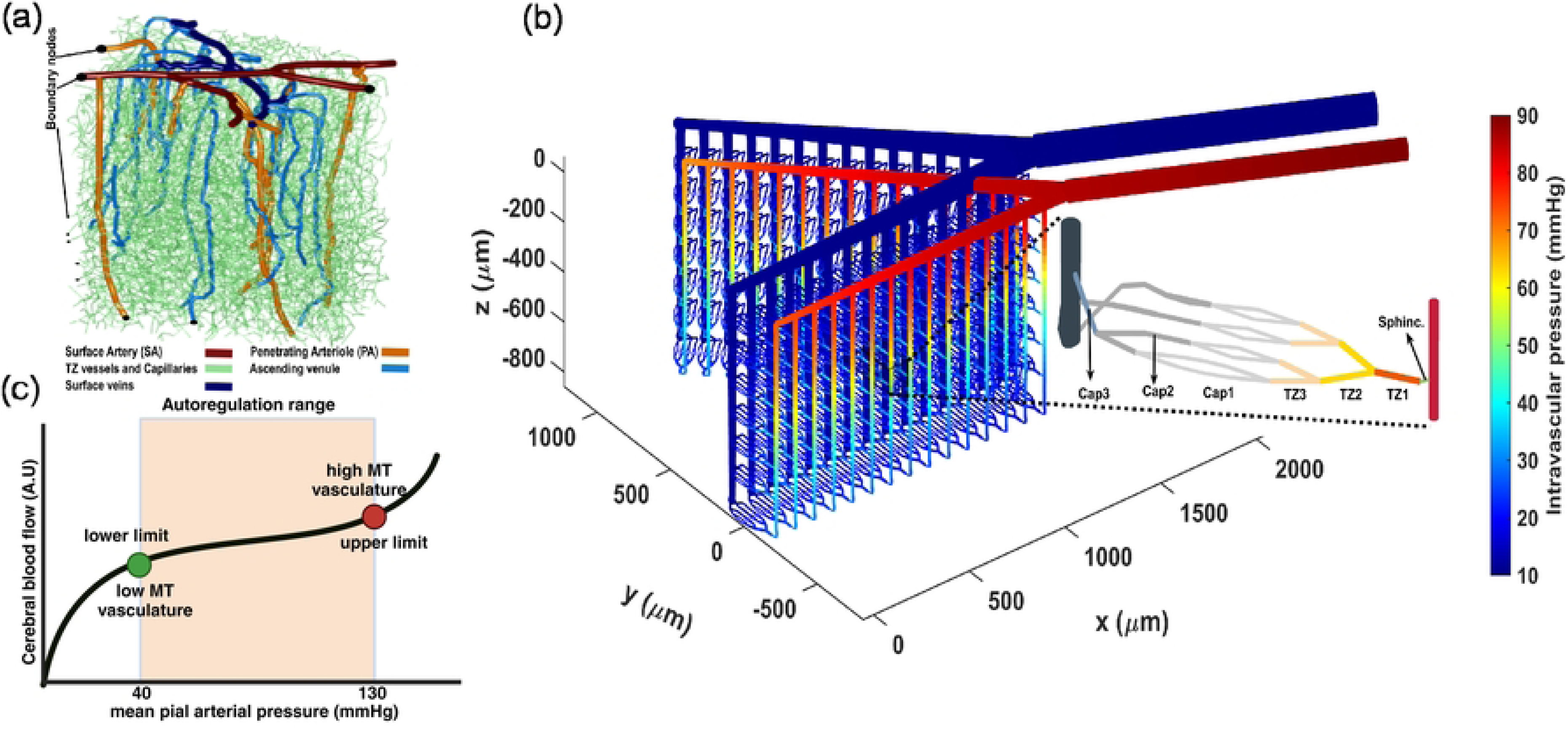
Overview of Cerebral Autoregulation Modeling in the Cerebrovascular Network. (a, b) segmented microvasculature structures:(a) is rendered from coarsely segmentation version of realistic in-silico mouse brain vasculature, provided by Kleinfeld’s laboratory[44], while (b) is an artificial vascular model. The vast number of boundary nodes in (a) pose a significant challenge in conducting precise hemodynamic analysis. Additionally, the inability to clearly identify different vascular zones, particularly TZ segments, makes it unsuitable for our study. These issues are addressed in (b) by incorporating only two boundary nodes and a monotonous structure. This architecture made it possible to differentiate between TZ segments and capillaries and to assign suitable mural cell to each segment. The zoomed-in area in (b) shows the structure of the microvessels, including sphincter, three layers of TZ segments (up to the 3rd order), and three layers of capillaries (beyond the 3rd order). Capillary indices are assigned based on the convergence morphology, indicating the blood volume accommodated by each segment. The color coding illustrates the pressure distribution across the vascular segments, with boundary node pressures set at 90 and 10 mmHg. The depicted in-silico vasculature model aims to replicate a final major bifurcation of the Middle Cerebral Artery (MCA). This structure was inspired by the detailed reconstructed pial arterial network of mouse cerebrovasculature in studies by Adams et al. and Epp et al. [45, 46]. (c) Illustrates the autoregulation curve. In our in-silico model, we defined the lower autoregulation limit at 40 mmHg, where mural cells induce minimal resistance to passive distension across all vessels. Conversely, at the arterial boundary node set to 130 mmHg, mural cells exert maximum resistance to passive distension, marking the upper limit in our model. This delineates the autoregulation range, within which mural cells dynamically adjust to maintain steady blood flow.

It is recognized that during brain’s vasculature development, the formation of vascular walls is regulated to obtain certain key attributes. These include the luminal diameter of vessels and the expression level of contractile elements encasing vessels. Estimating these variables was our first step in constructing the artificial cerebrovascular model. In tackling the challenge of defining the diameter of vessel without accounting for internal hemodynamics, we adopted a general approach for all segments. We started by identifying the diameter of vessels at their maximum dilation state defined as the point where the force exerted by mural cells balances out the passive distension during a stepwise increase in the IP of the vessel. Beyond this point, any further increase in the IP results in constriction since the force from mural cells overrides the passive distension and initiates vessel constriction. Theoretically, this state of dilation can be viewed as placing the vasculature at the lower limit of its autoregulation curve. As shown in Fig 1(c), the autoregulation curve illustrates the vasculature’s capacity to dynamically adjust vessel diameters, ensuring consistent blood perfusion in capillaries independent of the IP at the feeding points. Thus, the moment when constriction begins, due to an increased level of MT in vessels, can be considered theoretically as the lower limit of the autoregulation range. We began by analyzing images of murine cerebral vasculature to extract general cortical depth-dependent trends in vascular morphological features and incorporating this data into the modeled luminal vessel diameters. We specifically aimed to achieve uniform hemodynamics within capillaries when the IP in the main vasculature artery was set to 40 mmHg, which is a hypothetical value representing the lower limit of the autoregulation in our model. After achieving this uniformity, we introduced other factors, such as various expression levels of contractile elements in different segments of the vasculature. This approach enabled a systematic examination of cerebral autoregulation in our model.

#### A1: Cortical depth-dependent variations in cerebral vasculature morphology

Our analysis of the publicly available two-photon-imaged murine cerebral vascular data [47, 48] shows a cortical depth-dependency in morphological characteristics of the brain vasculature. Notably, aligned with Lauritzen et al.’s findings about sphincters [49, 50], we observed that these newly identified vascular segments typically form at strategic locations to help providing uniform hemodynamics across capillaries at various cortical depths. A sphincter, characterized by a bulbous dilation of the lumen following constriction, is usually located at the origination point of the transitional zone (TZ) - where arterioles transit to capillaries. Fig 2(a) show a PA that bifurcates into two daughter branches. A sphincter is visible at the origination point of the left branch in this figure. Our statistical analysis of sphincters, using the two-photon database, including the examination of 56 PAs across a cortical depth range of 0-600 µm in 12 mice (as detailed in Method), showed a close correlation between the likelihood of sphincter formation, cortical depth, and the ratio of daughter-to-parent vessel diameters. This result confirms the findings of Zambach et al.[49]. As depicted in Fig 2(b), our analysis revealed a noticeable decrease in the probability of sphincter formation at the bifurcation points as PAs delve deeper into the brain. Additionally, the results showed a strong correlation between the daughter-to-parent diameter ratio and the formation probability of sphincters, with sphincters being highly likely to form where the daughter branch diameter is less than that of the parent branch (Fig 2(b)). This feature is exemplified in Fig 2(a), where a sphincter is visible at the origination point of the left branch, with a lower daughter-to-parent diameter ratio, unlike the right branch where a sphincter is absent. Moreover, our analysis of the diameters of TZ segments in this dataset indicated a consistent enlargement trend in the average diameter of these vessel segments as PAs penetrate deeper into the cortical tissue, as shown in Fig 2(c).

**Fig 2.**
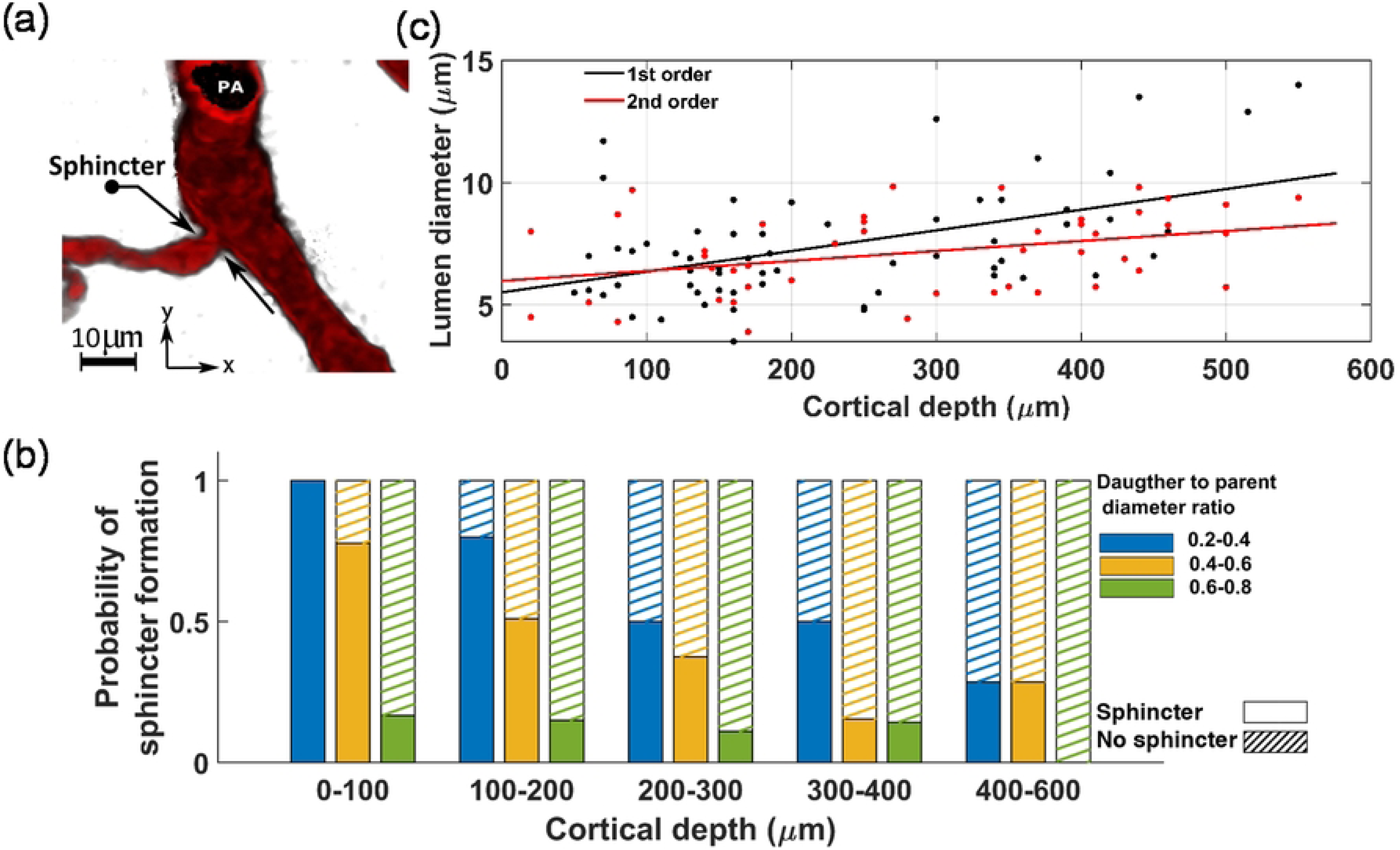
Quantitative analysis of sphincter density and TZ vessel diameters in mouse brain vasculature, (a) illustrates the presence of sphincters at PA-TZ bifurcation points. A sphincter is shown at the left bifurcation point, characterized by a low daughter-to-parent diameter ratio. In contrast, the right bifurcation point, with a larger daughter-to-parent diameter ratio, has no sphincter, (b) statistics of sphincter formation as a function of the daughter-to-parent diameter ratio at the PA bifurcation points across various cortical depths. The analysis highlights a close correlation between the cortical depth and the likelihood of sphincter formation at PA-TZ bifurcation points, (c) shows an increase in the diameter of TZ vessels as arterioles penetrate deeper into the brain tissue. Vessels are categorized by their order of bifurcation: the ‘1st-order’ TZ vessels emerge directly from the parent PA, while the ‘2nd-order’ TZ vessels are formed from the division of the 1st-order TZ vessels.

#### A2: Design of the in-silico vasculature at the maximum dilation point

The in-silico brain vasculature, shown in Fig 1(b), is a graph-based network composed of a series of interconnected nodes and segments. Each segment is modeled by a cylindrical tube with specific length and diameter that determine the resistance to blood flow according to Poiseuille’s law [51]. To improve the precision of the hemodynamic analysis, we designed this artificial vasculature with only two boundary nodes: a pial artery node for blood delivery and a vein node for blood drainage. This network features a pial (surface) artery that bifurcates into two branches, from which 15 PAs diverge and penetrate deep into the cortical tissue. We incorporated a total of 3 diverging bifurcations and up to 3 converging unifications connecting a feeding arteriole to a draining venule, drawing upon the findings of a study by Kleinfeld and colleagues[52]. Moreover, based on observations by Li et al.[9], blood flow in vessels that bifurcate from PAs was found to be nearly threefold larger than the flow in vessels on the venous side of the capillary bed, which direct blood towards ascending venules. Consequently, adhering to the principle of mass conservation, they proposed that the number of capillaries directing blood towards the venous side is approximately threefold larger than the number of branches carrying blood from arterioles to the capillary bed. A detailed view of the designed microvessels’ structure is provided in Fig1(b), illustrating a sequence beginning with a sphincter, followed by three diverging layers of TZ segments, and ending with three layers of converging capillary segments. The proposed morphological structure of capillary bed was designed to achieve smooth divergence and convergence and a good level of uniformity in vessel distribution, while adhering to a simple 2-to-1 convergence/divergence ratio. This results in eight non-bifurcating capillaries post TZ divergence that almost uniformly converge into three segments before connecting to an ascending venule.

In our in-silico vasculature, due to the monotonous structure of the capillary beds in all cortical depths, we assigned capillary branching order indices based on the blood flow level in each capillary segment. Accordingly, non-bifurcating capillary segments with the lowest blood flow were categorized as Cap1 segments. The first set of converging capillaries, handling blood flow twice that of the non-bifurcating capillaries, were designated as Cap2. Lastly, those capillaries accommodating a flow about four times higher than the non-bifurcating capillaries were classified as Cap3 segments. The length and diameter of various compartments within the network were established in accordance with recently published data showing the distribution of vessels’ characteristics across different vasculature zones[8, 53, 54]. The length of each arteriolar segment was approximated to match that of an arteriolar endothelial cell (EC), typically around 30-35 *µ*m[54, 55]. The lengths of sphincters were set to 4 *µ*m and each microvessel segment (TZ vessels and capillaries) to 44 *µ*m. This adoption of relatively long segment lengths (coarse segmentation) aimed to enhance the computational efficiency.

To establish homogeneous hemodynamics across capillaries at various cortical depths, we made several assumptions regarding the diameters of network segments. In line with our cause-and-effect assumption, we optimized the diameter of vessel segments at their maximum dilation to tune the blood flow in our vasculature within its lower limit of autoregulation curve. This optimization was aimed to ensure that, even with the IP in the main arterial node set within the lower physiological range, capillaries in all non-bifurcating branches have a hemodynamic with a Gaussian distribution, aligning with the range of values observed in in-vivo experiments and other computational studies.

Sphincters, TZ vessels and capillaries, classified as microvessels, account for a significant portion of the overall vascular resistance owing to their small internal diameter and the considerable distance that an RBC traverses within these segments. Thus, determining the diameter of these segments accurately is crucial. Several studies have measured microvessels’ diameters using in-vivo techniques[8, 56, 57, 58]. However, under normal conditions, vessels typically maintain a moderate level of MT, meaning that measured values do not reflect the diameter at the state of maximum dilation. Nevertheless, if we consider that a specific level of MT is typically present in vessels under realistic conditions in a certain mean arterial pressure, examining the diameters of vascular segments in in-vivo conditions can still provide valuable insights into their maximum dilation. In this regard, Watson et al. elucidated that TZ segments predominantly have a luminal diameter that is 20-25% larger than capillaries[8], aligned with the nomenclature we adopted in this article where TZ segments are the 1st to 3rd bifurcated vessel branch post-PA and capillaries are those beyond the 3rd order branch. Additionally, their research pointed out that the initial branch of TZ segments, TZ1, generally has a luminal diameter about 20% larger than that of the subsequent TZ segments (TZ2 and TZ3). In concordance with their statistical analysis and considering the larger contraction level in vessels in-vivo compared to their maximum dilation state, we estimated an additional 15% dilation on top of their reported values to represent the maximum dilation state of these micovessels more realistically. In particular, we set the TZ1 segments’ diameters based on a Gaussian distribution with mean value 5.5*µ*m and 0.25 standard deviation (*D*_TZ1_ ∼ *N* (5.5, 0.25^2^)), and subsequently, diameters of TZ2 and TZ3 segments were defined by (*D*_TZ2_ ∼ *N* (5, 0.25^2^)), and (*D*_TZ3_ ∼ *N* (4.5, 0.25^2^)). The diameters of Cap1 segments were also modeled by a Gaussian distribution (*D*_Cap1_∼ *N* (4, 0.25^2^))[8, 20, 34, 59]. For the two layers of converging capillaries, we assumed larger diameters (*D*_Cap2_ ∼ *N* (5.5, 0.5^2^)), and (*D*_Cap3_ ∼ *N* (6.5, 0.5^2^)). This adjustment accounts for the larger flow of RBCs in these segments, which potentially necessitates a larger cross-sectional area to accommodate the increased flow[56]. Additionally, blood pressure in these capillaries that are close to veins is typically low, which requires larger cross-sectional areas to provide low resistance to blood flow. The maximum dilation state of PAs was preliminarily estimated to be 18 *µ*m, which is slightly larger than the average basal diameter observed in in-vivo measurements made at the somatosensory cortex of mice that ranges between 10-15 *µ*m[14, 31, 60, 61, 62]. Also, diameter of pial artery segments was initially approximated to be around 60 *µ*m, a value derived from the physiological range observed in both in-vitro and in-vivo studies[62, 63, 64]. These initial estimates are later refined in this section to more accurately represent the maximum dilation state of these vessels.

To perform hemodynamic analysis within our designed vasculature, we set the IP boundary conditions to 40 mmHg for the pial artery node and 10 mmHg for the vein node. These values fall within the lower physiological range, aligning with our premise that our maximally dilated vasculature model would function optimally under low IP conditions closer to the lower end of the autoregulation curve. Hemodynamic analysis was then conducted, as detailed in Methods, to calculate blood flow, velocity, hematocrit, and pressure values across all segments of the artificial vasculature.

Multiple factors influence the variability of hemodynamic variables as observed in in-vivo studies [8, 9, 10, 11, 65] and computational analyses [19, 31, 32, 34, 35]. The key factors among these are the anatomical location of the vessel where the hemodynamic variables are measured, and the current state of the vasculature in its autoregulation range. To evaluate our in-silico vasculature’s ability to replicate actual conditions, and to mitigate the impact of the aforementioned factors causing hemodynamic variability for comparative analysis, selecting a comparatively stable physiological parameter was essential. We chose capillary blood flow as this key measure. Our rationale was based on the principle of autoregulation stating that irrespective of a capillary’s position within the brain’s vasculature, and under the condition of adequate blood oxygen and nutrients in experimental settings, blood should have a consistent range of flow throughout these vessels. Given the convergence of vessels within the capillary network, which can cause hemodynamic variability in capillaries [9, 11, 66, 67], we chose the blood flow in non-bifurcating segments, Cap1, as a representative measure of blood flow in capillaries. Considering their prevalence in the capillary bed, these segments constitute 56% of the overall vascular length of microvessels in our in-silico vasculature.

In in-vivo studies reported by Watson et al.[8] and Li et al.[9], RBC flux distribution in capillaries is typically a semi-Gaussian distribution centered around a mean value but with a skew towards higher fluxes[11, 66]. This tail in the distribution likely reflects the higher flow in converging capillaries, similar to what we modeled in our in-silico vasculature. Given that this pattern was observed in in-vivo experiments, it seems reasonable to consider the mean of this distribution as a desirable operating point of a healthy vasculature. A narrow distribution around this mean would ensure that most capillaries experience a similar blood flow, thereby minimizing instances of abnormally high or low flow. Computational studies typically report mean blood flow in capillaries ranging between 0.3 to 0.8 nl/min[19, 31, 32]. Correspondingly, RBC velocities measured in capillaries in in-vivo settings generally range from 0.4 mm/s to 1 mm/s[8, 9, 10, 11, 65]. Assuming single-file RBC movement in capillaries, it is reasonable to assume that the blood flow velocity in the centerline of these vessels approximates the RBC speed. Therefore, the measured RBC velocity in in-vivo settings, in a capillary with a typical diameter of 4 *µ*m (*D*) [8, 56, 68, 69], calculated by Eq. 1, results in a blood flow rate (*Q*) of approximately 0.3 to 0.75 nl/min. We selected a value near the median of this range, 0.5 nl/min, as the optimal blood flow rate for Cap1 segments in our in-silico model. Therefore, for an optimally designed in-silico vasculature, we anticipated that the blood flow within its non-bifurcating capillary segments (Cap1) to be a random variable with a narrow Gaussian distribution centered around 0.5 nl/min.

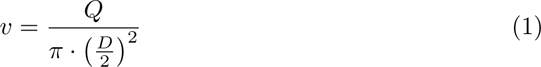

Preliminary hemodynamic simulations conducted using our artificial vasculature underscored the importance of incorporating cortical depth-dependency in vessel luminal diameters across different vasculature zones, particularly sphincters and TZ segments, as analyzed in Fig. 2. Neglecting this depth-dependency led to a disruption in the uniformity of RBC flux across capillaries at different cortical depths. Our results from these simulations under two distinct scenarios are presented in Fig. 3(a). In the first scenario, we strategically positioned sphincters at their optimal locations and modeled narrower diverging TZ vessels in upper layers. We adopted a simplified linear relationship between the sphincters and TZ vessels diameter (*D*_sphinc*_i_*/TZ*_i_*_) and cortical depth (*Z*). In our model, sphincter diameters ranged from 3 *µ*m in the superficial layer to 6.6 *µ*m in the deep cortex, computed using the equation *D*_sphinc*_i_*_ = 3 + 3.6 × cdr(*Z*_sphinc*_i_*_), where 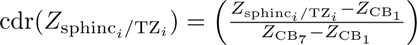 represents the cortical depth ratio. Here, *Z*_CB_ is the depth of the *m^th^* capillary bed (*Z*_CB1_ = 90*µm, Z*_CB7_ = 840*µm*), and (*Z*_sphinc*_i_*/TZ*_i_*_) is the depth of the capillary bed where the *i^th^* sphincter or TZ segment is located. For TZ vessels, a marked cortical depth-dependency in their diameters was observed, as shown in Fig. 2(c) (40-50% for a 400 *µ*m difference). This was modeled by a linear cortical depth-dependent scaling of the Gaussian distribution allocated to TZ segments, defined by *D*_TZ*_ni_*_ = *D*_TZ*_n_*_ × (1 + 0.8 cdr(*Z*_TZ*_ni_*_)), where *n* is the branch order. This leads to an approximate 80% increase in the diameter of TZ segments at 840 *µ*m depth compared to those at the depth of 90 *µ*m.

**Fig 3.**
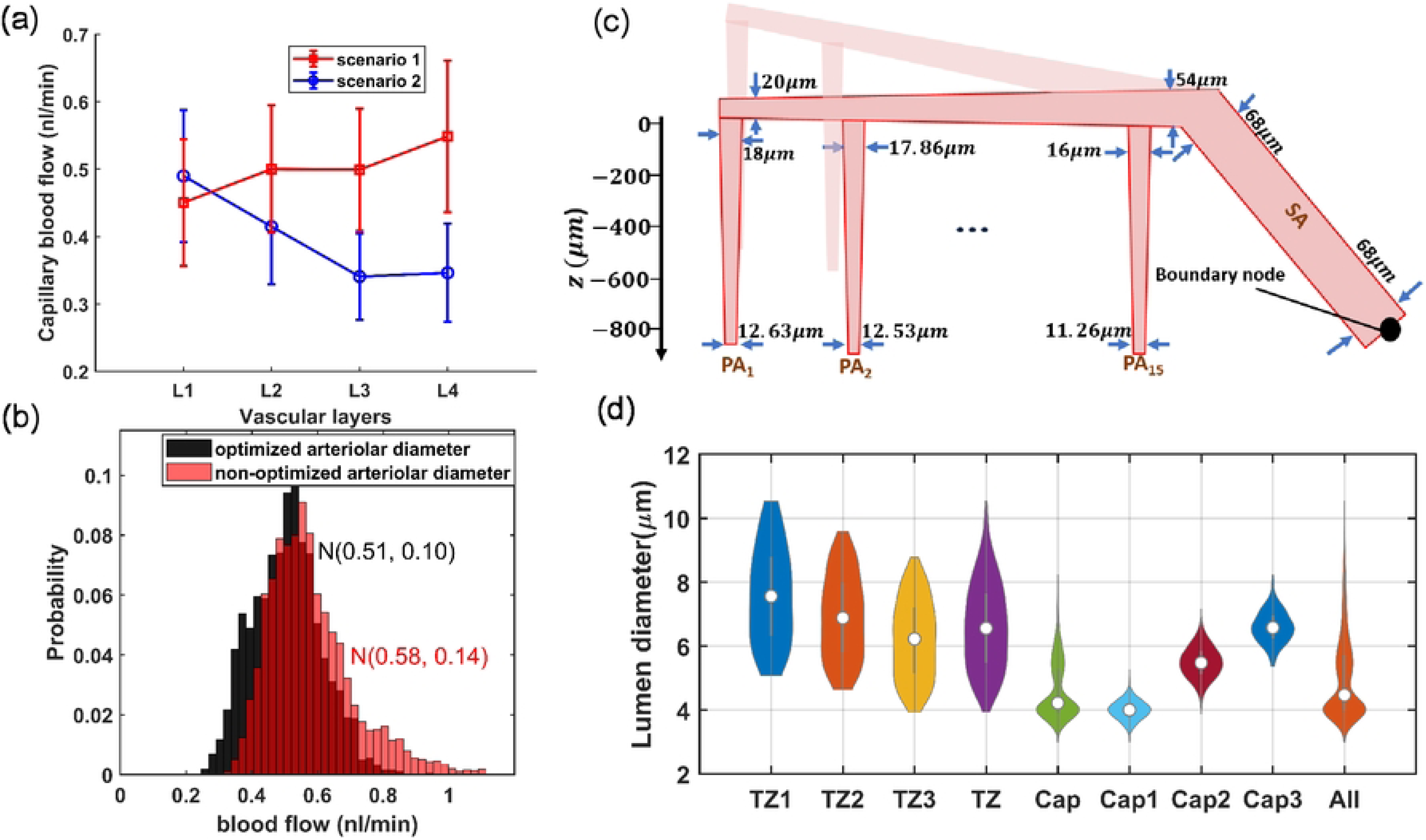
Design and optimization of cerebrovasculature model. (a) Blood flow in non-bifurcating capillaries across various cortical depths under two scenarios. Scenario 1 shows a more uniform distribution of blood flow, provided by the presence of sphincters and narrower superficial TZ segments, along with PAs that narrow as they penetrate deeper. In scenario 2, the absence of sphincters (equal 5 *µm* diameter for all cortical depths) and uniform diameters of TZ segments across all depths(*cdr*(*Z*) = 0), combined with a constant diameter of PAs (no narrowing along the penetration), result in a non-uniform distribution of blood flow across different cortical depths. (b) Presents a probability histogram of blood flow in Cap1 segments under two distinct scenarios. In the non-optimized scenario, all pial artery segment diameters were set to 60 *µm*, and PAs do not narrow as they approach the arterial boundary node. Non-optimized arteriolar diameter, results in a broader and more uneven distribution of blood flow in the capillaries. In this scenario, superficial capillaries located closer to the arterial boundary node have higher blood flow compared to those located further away. (c) A 2D schematic of the designed vasculature, emphasizing on the optimization of arteriolar diameters. In this optimization, we attempted to adhere to the principles of Murray’s law and took into account the proximity of some PAs to the arterial boundary node, where the IP is set to a fixed value. As a result, superficial segments of right PAs, being closer to higher pressure sources, are designed to be narrower, to adequately compensate for the elevated pressure levels. (d) Distribution of maximal dilation across microvessels in the designed vasculature. Cortical depth dependency considerations in TZ segments result in a higher standard deviation of diameters compared to capillaries. The ‘All’ category represents the diameter distribution of all microvessels in the model, where a noticeable peak is observed around 4 *µm* due to the larger prevalence of Cap1 in the microvasculature. The ‘TZ’ and ‘Cap’ groups denote vessels located within the transitional and capillary zones, respectively, without distinguishing between their specific indices.

Moreover, recognizing that deeper PA segments accommodate lower blood flow and consequently require smaller diameters than superficial segments, we applied Murray’s minimum-cost hypothesis as a guide to determine the rate at which each PA narrows. According to Murray’s law, the volumetric flow rate should be proportional to the cube of the vessel’s radius, implying that uniform wall shear stress (WSS) across the vascular network optimizes the system for minimum energy expenditure in fluid transport[38, 70]. While our design principle was shaped by insights from Murray’s hypothesis, we did not rigorously implement it in a mathematical sense. A precise application would entail a detailed mathematical optimization process, necessitating an accurate cost function and an advanced level of hemodynamic analysis, where WSS is determined based on equations accounting for the various blood velocity along the vessel axis. Our goal was to achieve a similar WSS across all segments, as formulated by Eq. 2, which provides a simplified equation for calculating WSS in a cylindrical vessel based on blood flow rate *Q*, apparent viscosity of the blood *µ*, and vessel diameter *D*. With this guiding principle, we formed PAs to lineally narrow from 18 *µ*m at the surface to 12.6 *µ*m in deeper layers. This gradual change in diameter was designed so that deepest segments of a PA in our vasculature (each PA segment feeding a single capillary bed (*Q*_PAdeepest_ = *Q*_Capillary bed_)), would have approximately the same WSS as the more superficial segments feeding seven capillary beds (*Q*_PAsurface_ = 7 *× Q*_Capillary bed_). Equalizing WSS in these segments requires that 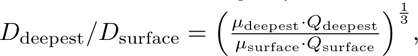 where the ratio 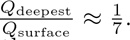 Due to the imprecise 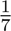 flow ratio in these segments as well as variations in the apparent viscosity of blood in microvessels with diameters ranging from 5 to 20 *µ*m[71], the ratio *D*_deepest_*/D*_surface_ = 0.7 yielded approximately identical WSS in both the deepest and surface segments of the PAs in our simulations. This is because the equation used to calculate blood viscosity and resistance of each segment in our hemodynamic analysis is based on Secomb and Pries studies [51, 71, 72], which proposed an empirically-derived estimation of the dependence of blood viscosity on the lumen diameter of the vessel segment.

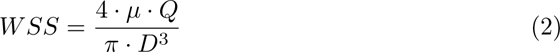

In scenario 1, PAs are narrowed with depth and cortical depth considerations are applied to sphincters and TZ vessels; however, these adaptations are not included in scenario 2. To analyze hemodynamics across different cortical depths, we categorized vascular layers differently from conventional cortical layer classifications, which are based on cellular composition and function at various depths. We defined four vascular layers within the cortex, L1-L4, each 210 µm thick with uniform capillary density. As Fig 3(a) shows, the first scenario achieves a uniform range of blood flow across non-bifurcating capillaries at various cortical depths, consistent with the 0.5 nl/min expected in capillaries. This uniformity indicates that the capillary blood flow distribution is closer to a narrow Gaussian distribution, a characteristic indicative of an evenly distributed energy supply throughout different brain areas which is well aligned with the conditions observed in realistic physiological settings [9]. Conversely, the second scenario leads to noticeable variations in blood flow across capillaries at different cortical depths, deviating from the expected flow in capillaries, particularly in deeper layers. These depth-dependent cerebrovascular morphological characteristics are essential for ensuring that the flow in deep capillary beds, with the lowest possible IP gradient at the lower limit of the autoregulation range, remains at levels comparable to those in superficial capillaries. Moreover, the progressive narrowing of PAs upon penetration plays a pivotal role in reducing flow in deep capillaries when deep TZs are wider, thereby tuning the pressure gradient and vascular resistance to equalize blood flow across capillaries at all cortical depths.

While ensuring uniform blood flow in capillaries connected to a specific PA across various cortical depths was crucial in our design, it was not the only criterion for adjusting vessel diameters in our network. Another aspect of our design was the gradual narrowing of arterioles as they approach to the arterial boundary node. This feature is essential to maintain consistent hemodynamics in capillaries, irrespective of their proximity to the main brain vasculature feeding point. To incorporate this aspect into our artificial vasculature, we posited that at the maximum dilation state, each PA’s surface segment would diminish by 0.14 *µm* as it approaches the arterial boundary node. Consequently, in our model, the diameter of the surface segments of the leftmost PA (PA1) was set to 18 *µm*, and the rightmost PA (PA15) to 16 *µm*. Note that due to the monotonous structure of the designed vasculature, where every PA feeds 7 similar capillary beds, this difference in the maximum diameter of PAs with a similar flow would result in varying WSS across different PAs (Eq 2), which is not in complete agreement with the Murray’s law. However, for simplicity, we did not reduce the number of capillary segments fed by PAs of smaller internal diameters. We also gradually narrowed pial arteriolar segments, as guided by Murray’s law, to define their maximum dilation states. Starting at 20 *µ*m diameter for pial segments adjacent to the leftmost PA (PA1)—a value close to the diameter of that PA’s superficial segments—we linearly increased the diameter to 54 *µ*m for segments following all 15 successive PA bifurcations from that daughter pial artery branch. This arteriolar diameter optimization strategy aimed to achieve approximately the same WSS in the leftmost segments of the daughter pial branch, feeding a PA as in the segments after PA15, which feed 15 PAs (Eq 2). Given the similar structural configuration and resultant blood flow at the origination point of the two daughter pial branche<u>s</u>, we multiplied the cube root of 2 to the diameter of these initial segments 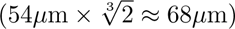 to determine the diameter for the terminal segment of the parent pial artery before its bifurcation into daughter branches. Since this parent pial artery experiences no further bifurcations, and thus no changes in blood flow along its path, the segments’ diameter remains constant up to the pial boundary node. These design considerations ensured that the distribution of blood flow in non-bifurcating capillary segments throughout the in-silico vasculature closely aligns with a Gaussian distribution. Fig 3(b) displays the probability histogram of blood flow in non-bifurcating capillary segments (Cap1). This figure illustrates that networks with optimized arteriolar diameters achieve a distribution of blood flow that is consistent with a Gaussian distribution. Conversely, networks with non-optimized arteriolar diameters demonstrate a more uneven distribution of blood flow. Fig 3(c) illustrates arterioles at their optimized maximum dilation state, and Fig 3(d) shows the distribution of the adjusted maximum dilation state across microvessels, all modeled under an artery boundary pressure of 40 mmHg (30 mmHg pressure gradient in the vascular network).

In designing our cerebrovascular model, we noticed the discrepancy that exists in studies regarding the ratio of number of PAs to the number of ascending venules (AVs), which typically ranges from 1-3 AVs for every PA in rodent animals[73, 74, 75, 76]. This discrepancy suggests that, in reality, AVs carry a lower volume of blood compared to PAs, despite having basal diameters that are generally comparable[44, 77, 78, 79]. To account for this in our model—where an equal number of PAs and AVs were included for simplicity—we assigned larger diameters to AVs than PAs. This design choice ensures minimum IP drop as blood moves through venular segments, aligning with findings of previous researches[31, 32, 34]. Consequently, diameters for AVs were set to 30 *µm* for surface segments and 20 *µm* for deep segments[80]. To further prevent IP drop along the pial vein, diameters of these segments were also set to larger values than those of the pial arteries, progressively increasing from 30 to 90 *µm* from the leftmost to the rightmost segment. While this simplification might seem significant, it does not impact the investigation of cerebral autoregulation critically, as the model ensures negligible IP drop occurrence in venous segments[25], maintaining the focus on arterial segments’ roles.

#### A3: Analysis of in-silico vasculature at the maximum dilation point

As shown in Fig 4(a-b), blood flow and flow velocity in non-bifurcating capillaries are at lowest values compared to other branching indices. Blood flow is evenly distributed in Cap1 segments; therefore, at every convergence following Cap1, blood flow doubles. Moreover, our analysis suggests that not distinguishing between converging and non-converging capillary segments can broaden the Gaussian distribution of blood flow, leading to a distribution which has a skew towards larger values. This effect, primarily linked to converging capillary segments, subtly alters the overall expected distribution of RBC flux, which is typically dominated by the more abundant Cap1 segments. Consequently, this leads to a noticeable tailing effect in the distribution patterns of RBC flux within the capillary networks, as observed in multiple in-vivo studies[9, 11, 66]. Furthermore, Fig 4(c) illustrates the relationship between the diameter of capillaries and their blood flow velocity in the cerebrovasculature model. As shown, despite noticeable variations in both blood flow (Fig 4(a)) and the diameter of capillaries (Fig 3(d)), the blood flow velocity within capillaries consistently ranges from 0.5 to 1 mm/s. This finding aligns with results from previous in-vivo studies [8, 81, 82, 83] in which lack of strong correlation between capillary diameter and RBC velocity was also observed. Such a fundamental morphological-hemodynamic relationship serves as a determining validation measure for the design of our in-silico vasculature, particularly in terms of the morphological characteristics that we incorporated in the design of the capillary networks. As illustrated in the green violin plot of Fig 4(b), blood flow velocity across all capillaries is relatively uniform with a narrow Gaussian distribution that aligns with in-vivo observations [19, 66, 82]. This distribution lacks the apparent tailing effect observed in the blood flow distribution (green violin plot in Fig 4(a)), suggesting a more uniform RBC velocity across all capillaries.

**Fig 4.**
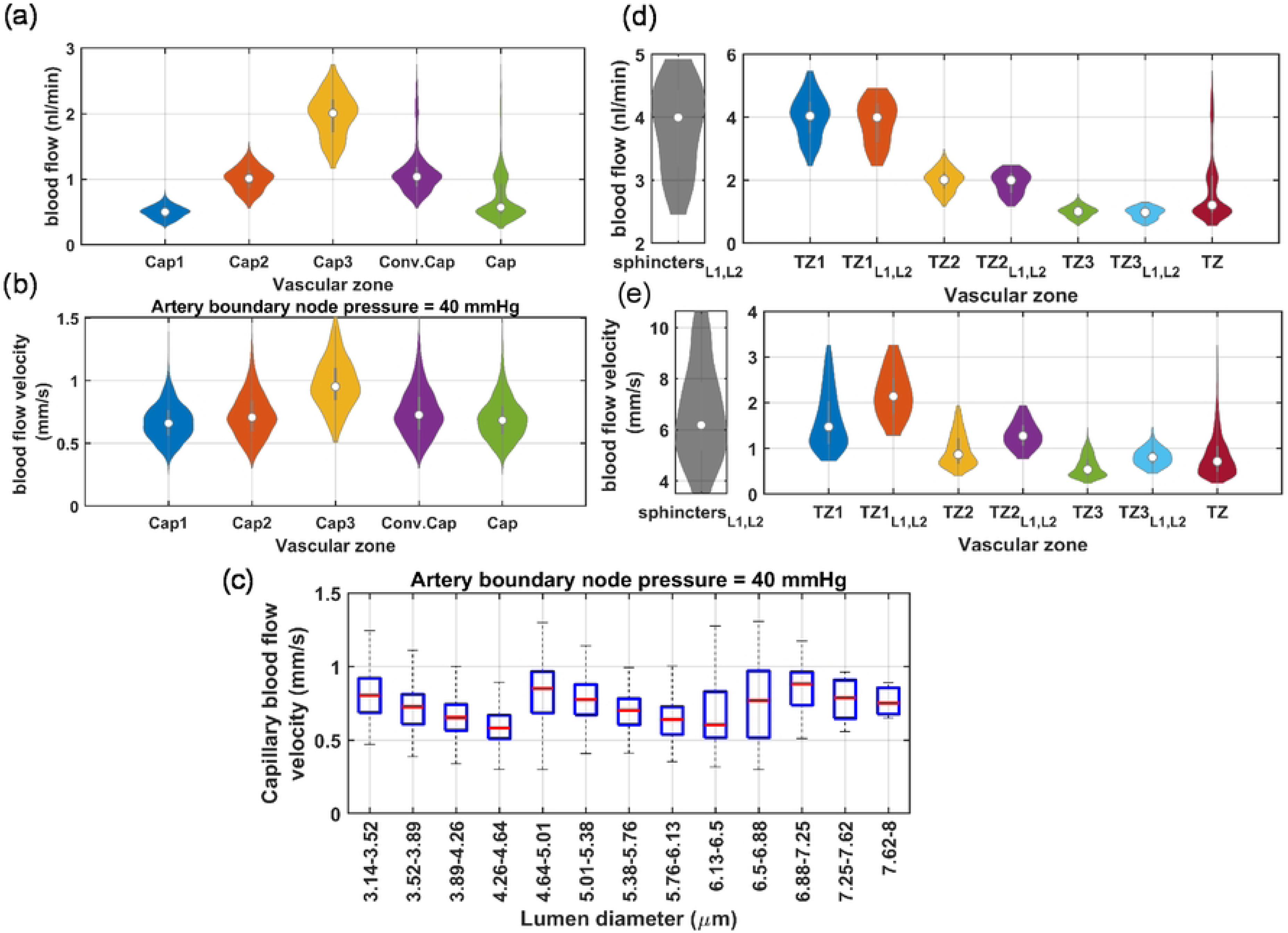
Analysis of hemodynamics in microvessels of the cerebrovascular model at maximum dilation. (a) Shows that blood flow in capillaries is predominantly centered around the characteristics of Cap1 due to their larger prevalence in the network. The distribution tail is associated with the converging capillaries, (b) in contrast, the blood flow velocity across all capillaries is relatively uniform and has a narrow Gaussian distribution, as displayed by the green violin plot, (c) shows the relationship between the luminal diameter of capillaries and their blood flow velocity. This data shows no strong correlation in this relationship; however, three distinct decremental trends are observable, each associated with a different capillary index. This observation is consistent with the assignment of capillary indices and their luminal diameters, which were determined based on the typical range of blood flow that they accommodate. Within each capillary index category, larger diameter capillaries have lower blood flow velocity, (d-e) shows blood flow and blood flow velocity in TZ segments, highlighting noticeable variations across different branching orders and cortical depths. It is important to note that the blood flow velocity represented here corresponds to an arterial boundary node pressure of 40 mmHg, which is the point where vessels are at their maximum dilation state. These values can be up to twice as high under elevated pressure, which is a topic that will be further discussed in Sec A5. The subscript “L1, L2” corresponds to vascular segments located at cortical depths from 0 to 420 *µm*.

In our analysis of the TZ segments’ hemodynamics, as displayed in Fig 4(d), depth-dependent variations in the luminal diameter of TZ vessels, results in variable resistance at the PAs’ bifurcation points. However, the combination of a larger resistance in capillary beds of upper layers and a higher IP gradient, as well as smaller resistance and a lower IP gradient in deeper layers, ultimately leads to a nearly equal amount of blood flowing into each capillary bed. Upon entry into the capillary bed, the blood flow is approximately halved at each successive bifurcation point within the TZ. Therefore, taking into account the three divergences before reaching the Cap1 segments, the blood flow in the sphincters and TZ1 segments is about eight times larger than the flow in the non-bifurcating capillaries.

While blood flow generally remains consistent across the similar branching order of TZ segments in our network, we observed some variations in the blood flow velocity within these segments, particularly in relation to the cortical depth. These variations are attributed to the depth-dependent characteristics of the sphincters and TZ segments in our model. Specifically, superficial TZ vessels, which are characterized by narrower maximum dilation states, typically experience higher blood flow velocity. Moreover, a decreasing trend was incorporated in the diameter of successive TZ branching orders in the cerebrovasculature model [14, 60] (Fig 3(d)), resulting in blood flow velocity diminishing to less than 50% after passing each bifurcation point. This trend is consistent with the RBC velocity measurements in TZ segments reported in studies such as those by Watson et al.[8]. However, a comprehensive analysis of blood flow velocity in TZ segments necessitates accounting for cortical depth variations. As shown in Fig 4(e), blood velocity in cortical depths between 0-420 *µ*m (L1-L2 in Fig 4(d-e)) is at the higher end of the blood flow velocity range across all branching orders. Furthermore, the prevalence of TZ3 segments, being four times more abundant than TZ1, significantly influences the average RBC velocity and flux in the TZ group (as shown in Fig 4(d-e)), tending to reflect the characteristics typical of TZ3 segments. This branching order and depth-dependencies are essential factors that need to be included in the analysis for accurate interpretation of the hemodynamic behavior within the transitional zone.

Hemodynamics in TZ segments is influenced not only by the cortical depth but also by the IP values at the vasculature feeding points, which determines the extent to which vessels are constricted relative to their maximum dilation state. Similarly, hemodynamics in PAs is also a function of IP variations at these feeding points. To explore the hemodynamic range across other vascular zones more effectively, it is essential to first refine the model to make it operational for artery boundary node pressures (ABNP) above 40 mmHg.

#### A4: Design of the in-silico vasculature model within the autoregulation range

Previously, we assumed that at the lower end of the autoregulation range, under the ABNP of 40 mmHg, the maximum luminal diameter of vascular segments is the only variable responsible for the uniform distribution of blood in different regions (Fig 3(a)). We assumed that by increasing the ABNP in a network of elastic vessels from 0 to 40 mmHg, all vessels would dilate and once the ABNP reaches 40 mmHg, the IP is distributed such that all vessel segments are at the state of maximum dilation and all regions are nearly fed equally. In this state, pial arteriolar segments, diameter of ∼68*µ*m, achieve their maximum dilation around 40 mmHg IP. However, because of the IP drop along the blood flow pathway, maximum dilation in deeper PA segments, diameters ∼11 - 13*µ*m, occurs at IP values around 30-35 mmHg. This variation indicates that the threshold force which initiates vessel constriction depends on the vessels’ luminal diameter and its intensity changes as a function of the position of the vessel in the vascular network. Therefore, based on this assumption, increasing ABNP above 40 mmHg triggers concurrent constriction in all vessels encased by mural cells.

To model vessel constriction at ABNP values larger than 40 mmHg, we need detailed information about the transfer function of individual vascular segments, which is a function that relates a broad range of their internal hemodynamics to their luminal diameter. However, obtaining these transfer functions for every vascular segment is technically infeasible. Therefore, we opted for a generalized approach to avoid determining segment-specific transfer functions. The goal behind the development of each vascular segment’s transfer function is to regulate blood flow throughout the network independent of the ABNP value. This means that for ABNP values larger than 40 mmHg, mural cells within each vascular segment should detect hemodynamics to induce an appropriate level of constriction force to maintain capillary blood flow at levels comparable to those at 40 mmHg. In our approach, we introduced the hemodynamic regulation constriction transfer function (HRCTF), to estimate this constriction force between the ABNP values of 40 to 130 mmHg, which is the hypothetical autoregulation range of our model. HRCTF is a function that indicates vessels constriction percentage relative to their maximum dilation to model autoregulation in the vasculature. However, due to variations in the expression level of contractile elements across different segments, with SMCs in PAs and SAs featuring larger constriction capacity compared to thin-strand pericytes in capillaries[53, 84], the constriction level is not uniform in all vascular segments. Therefore, in addition to HRCTF, we also introduced the relative contractility (RC) for each segment, which serves as an index to indicate the ability of a vessel segment to actively constrict and counteract the passive distention in response to ABNP changes. The product of the vasculature HRCTF and the RC indices of vascular segments estimates the level of constriction in all segments relative to their maximum dilation, also known as the percent tone of vascular segments. By incorporating these estimated percent tone values into the diameters of vascular segments at different ABNP values, we achieve a flat ABNP-flow relationship in our vasculature, to model autoregulation.

Before tuning the vasculature HRCTF for different ABNP values, first we should assign RC indices to each segment. The RC of each vascular segment denotes the potential of mural cells encasing that segment, compared to other segments, to exert force and contract the vessel. This parameter is associated with the relative expression level of contractile proteins in those cells. Some vascular segments, such as sphincters, produce larger pressure gradients over small areas. To withstand a large IP without rupturing, these segments are usually fortified by a larger concentration of fibroblast-synthesized collagen[85], and an abundance of contractile elements, particularly the *α*-SMA. Zambach et al. reported that, on average, the *α*-SMA expression in sphincters surpasses the amount found in the SMCs of PAs [49]. Our simulations indicated that sphincters, especially in the superficial layer, experience higher IP at their entry points, compared to the average IP in PAs, as displayed in Fig 5(a). This cortical depth-dependency suggests that a uniform RC index cannot be assigned to all sphincters or to all vascular segments located at different cortical depths. The lack of quantitative studies comparing expression levels of contractile elements in different mural cells across various cortical depths necessitates devising a method for allocating an appropriate RC index to each segment in the model.

**Fig 5.**
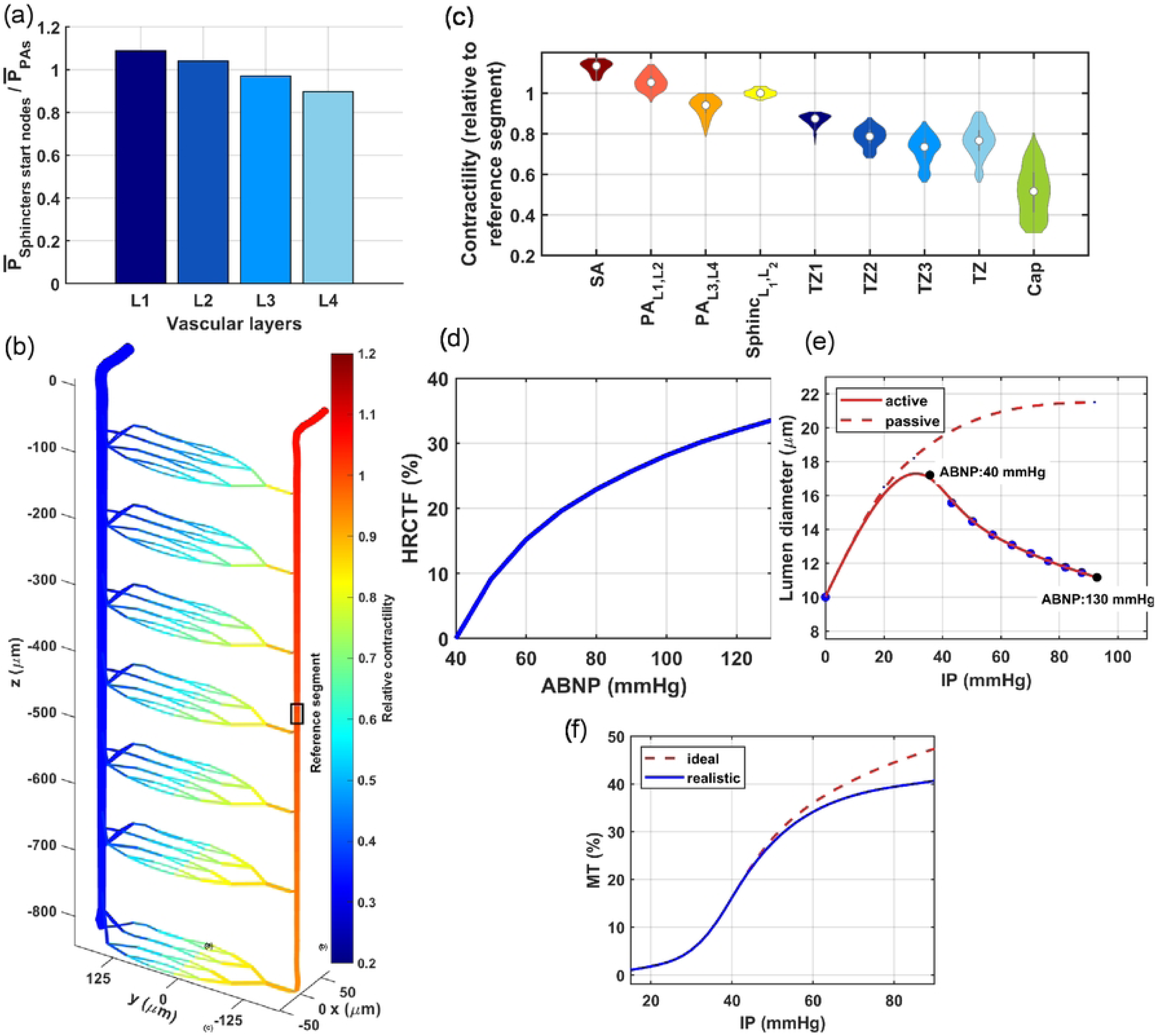
Designing a cerebrovascular model within the autoregulation range. (a-c) Relative contractility in the in-silico vasculature: (a) superficial sphincters typically experience higher average IP at their entry points 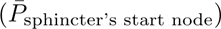 compared to deeper sphincters. This graph highlights the cortical depth-dependency in the contractility of different vascular zones, (b) illustrates the estimated relative contractility of vascular segments across a portion of the in-silico vasculature model. RC indices were estimated by dividing the IP of each segment to the IP of a designated reference segment in the maximally dilated vasculature, (c) statistical analysis of RC in the cerebrovascular model shows that superficial PA segments and sphincters possess larger contractility compared to deeper PA segments. TZ segments also have large contractility in contrast to capillaries which have the least contractility compared to preceding vascular zones. L1-L4 represent the four vascular layers, where each is 210 *µ*m thick in the model, (d) the non-linear characteristic of the HRCTF required to maintain constant mean capillary blood flow as a function of ABNP values. This relationship indicates the need for larger initial constriction, which lessens as the ABNP increases, (e) the solid line shows constriction in a PA segment at 100 *µ*m cortical depth. The dashed line represents the estimated passive distention of this segment when SMCs are inactive, (f) the solid line shows the MT of this segment based on the curves plotted in panel (e), calculated from 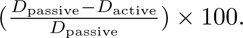 The dashed line illustrates the realistic MT profile for this segment, approximated by averaging the MT profiles of male and female mouse PAs as reported by Jeffrey et al.[86].

When ABNP increases from 40 mmHg, the force exerted by mural cells on the external surface of the vessel lumen must exceed the force exerted by IP on the internal surface to initiate constriction. Consequently, segments experiencing higher IP at 40 mmHg ABNP, such as SAs and superficial PA segments, need to generate stronger forces to counteract the IP force. Conversely, deeper PA segments do not require a substantial myogenic response since the IP is attenuated along the blood flow pathway. This variation in the IP force across different segments enabled us to estimate the relative contractility of each segment in our model by comparing the IP forces at 40 mmHg ABNP. Therefore, we determined RC values by normalizing the IP of each segment to a reference value, which was the IP of the middle segment in PA1. In this analysis, the assigned RC of the reference segment is 1.0. Segments with higher IP than the reference have RC values larger than 1.0, while the downstream segments from the reference have smaller RC values.

The graph in Fig 5(b) illustrates the distribution of estimated relative contractility of vascular segments in a portion of the in-silico vasculature model. As shown, superficial segments of PA have slightly larger contractility compared to deeper segments. While no study has explicitly reported a decreasing trend in *α*-SMA expression with the progression of PAs through the brain tissue, immunohistochemical staining for *α*-SMA in a slice of mouse cerebral vasculature, as shown in Hartman’s study, suggests that such a trend exists (Figure 1(b,d) of [53]). Furthermore, as illustrated in Fig 5(b, c), our analysis shows relatively uniform contractility in similar branch orders of TZ vessels across various cortical depths, albeit with marginally smaller values in superficial layers. In our vasculature design approach, we optimized lumen diameters to ensure a uniform distribution of blood flow across various cortical depths at 40 mmHg ABNP. Due to the need for superficial layers to have narrower sphincters and TZ vessels in their maximum dilation state to prevent excessive flow in their capillaries (Fig 3(a)), superficial TZ vessels experience low IP at the 40 mmHg ABNP level; consequently, the calculated RC indices for superficial TZ vessels are small. To the best of our knowledge, no study has investigated the expression level of contractile proteins in TZ vessels across different cortical depths. Nevertheless, Zambach et al. showed that when a sphincter is present at the PA bifurcation point, primarily in superficial layers (Fig 2), the expression level of contractile proteins in higher order branches of TZ vessels is noticeably small [49]. Fig 5(c) depicts the statistical analysis of RC across various vasculature zones. As shown, the RC of capillaries is on average 67% of that in TZ vessels. This is supported by the findings of Hartman’s study, which showed that constriction in cerebral capillaries following the transition from hypercapnia to normocapnia is about 60% of the constriction in TZ vessels [53]. Furthermore, Klug et al. showed that constriction in retinal capillaries is, on average, 57% of that in TZ vessels when the ABNP of the retinal vasculature was gradually increased [87].

After obtaining estimates of each segment’s RC index, we modeled the autoregulation in the vasculature. To accomplish this, we incrementally increased the ABNP from 40 to 130 mmHg in 10 mmHg steps. At each step, we calculated the HRCTF value such that, when multiplied by each segment’s RC index, the resulted constriction in vessels can maintain the average capillary blood flow at the level obtained at 40 mmHg ABNP (Fig 3(a)). RC indices are relative values, and the reference segment could be anywhere in the vasculature. Although the selection of a reference segment changes RC indices, multiplying these RC indices by the HRCTF derived from the same set ultimately yields the same percent tone estimates for all vascular segments and across all ABNP values. Moreover, we kept the vein boundary node pressure (VBNP) constant at 10 mmHg in our simulations as ABNP increased from 40 to 130 mmHg. We posited this because the majority of the pressure drop in vessels should occur in the arterial system before reaching capillaries and venous segments, which are encased by mural cells with minimum contractility and are unable to handle high pressure.

Fig 5(d) displays the HRCTF of our vascular model, or the percent tone of the reference segment, across various ABNP values within the autoregulation range. Fig 5(e-solid line) shows changes in the diameter of a PA segment located at 100 *µ*m depth with a relative contractility of 1.05. Based on our generalization approach, we assumed that vessels behave like elastic elements and begin to inflate from zero ABNP until they reach a threshold beyond which their muscles induce constriction. In our model, this threshold occurs for all vascular segments at 40 mmHg ABNP. The superficial PA segment depicted in Fig 5(e), achieves its maximum dilation approximately between 28-35 mmHg IP, aligning with observations of in-vitro pressure myography studies of PAs of mice cerebral vasculature with similar maximum luminal diameters [86]. The total constriction experienced by this segment throughout the entire autoregulation range is around a maximum of 35%, HRCTF (ABNP=130) × RC(PA_100_ *_µ_*_m_) ≈ 33.5% × 1.05, which falls within the reasonable range for maximum constriction that an arteriolar segment can undergo over the entire autoregulation range, as observed by Klug et al.[87].

The non-linear nature of the HRCTF indicates that to achieve autoregulatory plateau (flat ABNP-flow relationship) in a vascular network as ABNP linearly increases, vessels must constrict in a non-linear manner. This non-linear constriction is essential to provide a proportionately larger resistance to blood flow in the vasculature network for higher ABNP values, to ensure the stability of capillary blood flow, or equivalently, artery boundary node flow. This concept is shown in Fig 5(e). The figure shows that vessels initially constrict steeply. As the pressure further increases, even a small constriction is enough to generate a larger resistance. This behavior correlates with the findings from in-vitro studies investigating MT in isolated vessels, where arterioles constrict more sharply in response to initial step-wise pressure increases, followed by a gradual decrease in the constriction slope as the IP increases [86, 88, 89]. This pattern reflects the interplay between the passive vessel distention and the active force exerted by mural cells. Initially, mural cells can easily counteract the passive distention, but as the IP rises, inducing further constriction becomes increasingly challenging.

MT signifies the ability of mural cells to counteract passive distention, while the percent tone we estimated, indicates the level of vessel constriction necessary for flat autoregulation. To assess how closely the estimated percent tone matches realistic conditions, we need to estimate the MT of vessels, a parameter frequently reported in the studies of mouse cerebral vasculature. Estimating MT involves determining how vessels passively distend in response to an increase in IP when their muscles are inactive which reflects the vessels’ distensibility. Given the delicate nature of PAs with a luminal diameter between 10-20 µm, only the study by Jeffrey et al. has investigated MT in mouse cerebral PAs within this diameter range[86]. Drawing on data from Jeffrey’s study, the dashed curve in Fig 5(e) presents an estimated profile for the passive distention of a PA segment at 100 µm cortical depth. Jeffrey et al. showed that, similar to pial arteries[90, 91], MT in PAs is influenced by the basal activity of endothelial nitric oxide synthase (eNOS). Fig 5(f-dashed line) shows a representative MT profile from Jeffrey’s study. The dashed curve in Fig 5(f), derived from plots in Fig 5(e), shows the MT profile that results in flat autoregulation. Comparing the ideal MT profile with its realistic representation in Fig 5(f) suggests that the capacity of mural cells in mice cerebral arterioles might not support flat autoregulation throughout the entire range, potentially showing a slight slope in the flow-pressure relationship at the mid to upper autoregulation range.

#### A5: Analysis of in-Silico vasculature model within the autoregulation range

Fig. 6(a) displays the mean and standard deviation of blood flow in non-bifurcating capillaries across layers L1-L4 for different ABNP values. This vasculature was modeled to autoregulate blood flow by adjusting diameters of segments based on their estimated percent tone calculated by multiplying the HRCTF and RC indices. Simulation results show that in L1, and at the lower limit of the autoregulation range, the existence of narrower sphincters and TZ segments lead to a diminished capacity for low IP in the PAs to push RBCs through these narrow pathways. Consequently, blood flow in L1 is typically at its minimum at low ABNP. As the ABNP, and subsequently IP in the PA segments of L1 increase, the force driving RBCs through these narrow pathways also rises, which leads to an increased blood flow in this layer. The HRCTF was tuned to maintain constant mean capillary blood flow across all cortical depths. As a result, the inflow to the vasculature was kept constant. Therefore, while average blood flow in superficial capillaries (L1-L2) increases with rising ABNP, it decreases in deeper capillaries(L3-L4).

**Fig 6.**
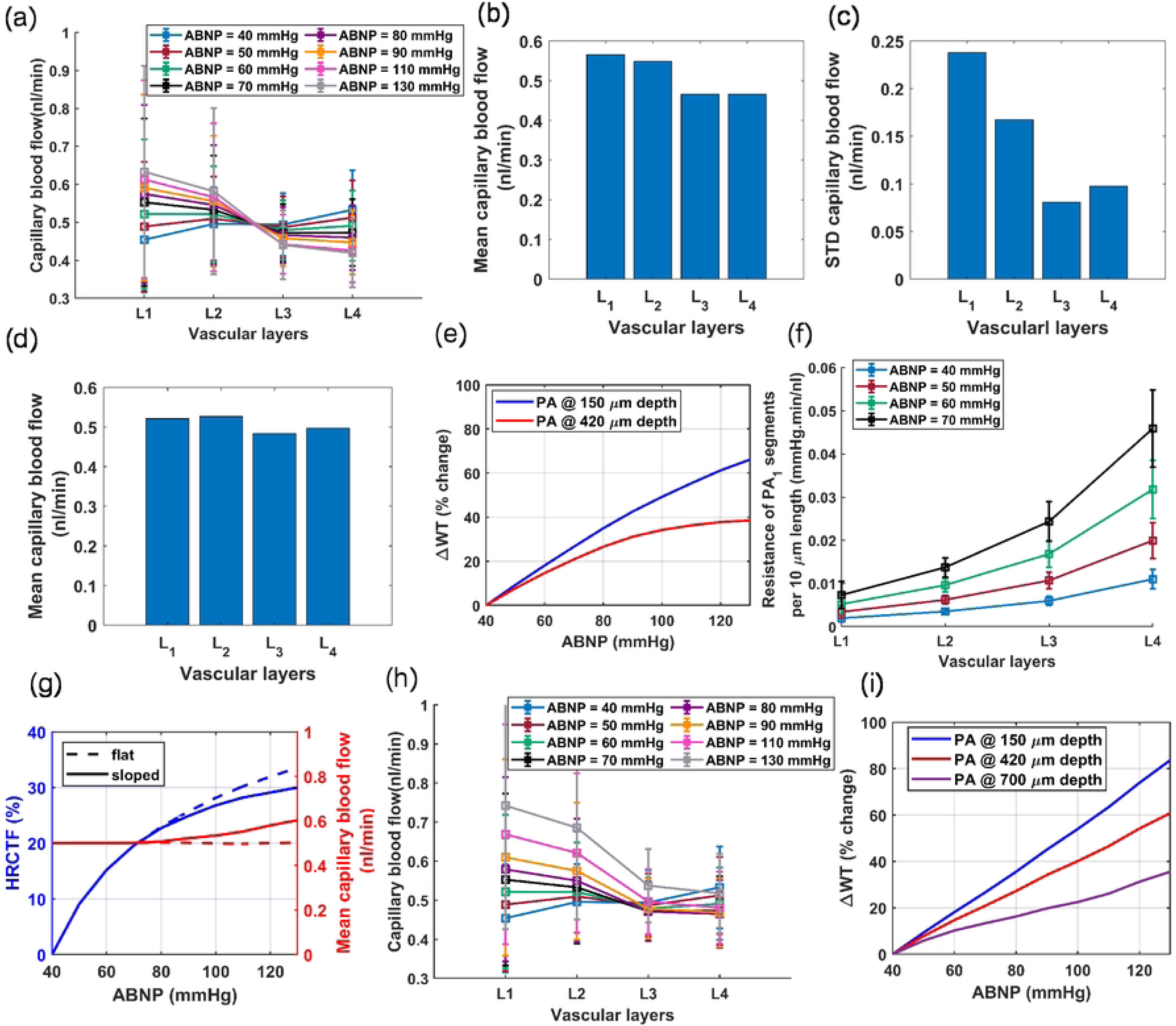
Validation and predictive accuracy of the cerebrovascular model, (a-f) dependency of blood flow in capillaries on the cortical depth and ABNP values: (a) mean and standard deviation of blood flow in Cap1 segments for various ABNP values, (b) layer-specific average blood flow in capillaries across the entire monophasic flat autoregulation range, (c) indicates that the superficial layers have a more nonuniform hemodynamic distribution in capillaries, (d) mean capillary blood flow across different vascular layers at physiological ABNP values (50-70 mmHg), (e) WT changes in superficial PA segments are more significant than in deeper segments across the autoregulation range. This suggests that superficial PAs are prone to larger distortions in their lumen diameters, since their MT mechanism must perform over a wider range of WT, (f) mean and standard deviation of blood flow resistance per 10 micrometer length of PA1. A larger variability in resistance is observed in L4 compared to upper layers. (g) The ideal and realistic vasculature HRCTFs, correspond to monophasic and biphasic autoregulation, respectively. (h) The mean and standard deviation of blood flow in Cap1 segments for various ABNP values when vessel lumen diameters were adjusted based on the biphasic HRCTF. (i) WT in vessels increases linearly when ABNP increases linearly, if the vasculature is modeled for biphasic autoregulation. This observation suggests that the myogenic response is potentially linearly potentiated with increasing WT; however, the decreased constriction ability of muscles in the sloped phase, is proven to be advantageous for the vasculature, as it prevents reduced blood flow in deeper layers at high ABNP values.

Next, we combined blood flow data from non-bifurcating capillaries at all tested ABNP levels (10 steps) into a single dataset and calculated the means and standard deviations across layers L1-L4, Fig. 6(b-c). Results of this statistical analysis of blood flow across different layers align well with Li et al.’s measurements of capillary RBC flux in layers I-V in the whisker barrel cortex of awake mice[9]. Li et al. observed that RBC flux in Layers IV and V is lower than in layers I-III. In our categorization of vascular layers, cortical layers IV and V approximately correspond to layer L3. As shown in Fig. 6(a), almost throughout the entire autoregulation range, capillaries in L3 consistently have lower blood flow compared to upper layers. Li et al. proposed that the increase in the capillary density in layers IV and V might be the reason for the reduced RBC flux in these layers. However, the lower blood flow in L3 of our vasculature model, where capillary density is uniform across all cortical depths, has a different cause.

Blood flow in a capillary bed is determined by the pressure gradient between the PAs’ bifurcation point and the capillary converging points leading to ascending venules, and the resistance to blood flow determined by the microvessels’ lumen diameter. IP is a systemic parameter, that is influenced by both the upstream and downstream segments and serves as the main factor in adjusting vessel lumen diameters through the MT mechanism. This means that to achieve uniform blood flow in all vascular segments throughout the broad range of ABNP, there must always be a balance between IP distribution and lumen diameters across the vascular network. In theory, achieving this balance across the entire autoregulation range seems impossible, especially given the relatively static nature of the passive vessel lumen diameter. The layered architecture of the brain vasculature suggests that, under the best-case scenario, the vasculature development regulatory mechanisms could only ensure a uniform blood flow distribution across all cortical depths within a narrow band of the autoregulation range (ABNP 50 mmHg in Fig. 6(a)). As depicted in Fig. 6(a-blue), at the maximum dilation state, capillaries in L1 have the least blood flow while capillaries in L4 have the largest, both slightly deviating from the ideal capillary blood flow of 0.5 nl/min. As the ABNP increases from 40 mmHg to 130 mmHg and with the potentiation of MT in vessels (Fig. 6(a)), blood flow in L1 increases and in L4 decreases, moving them closer to the ideal level, while L2 and L3 begin to diverge from this ideal value. At the mid to larger values of the autoregulation range (ABNP 80-130 mmHg), the potentiation of MT in deeper PA segments, which possess smaller maximum lumen diameters, could undesirably reduce the IP along the penetration below the preferred level. In our vasculature model, this IP reduction in deep PA segments occurs since the model was designed to function optimally within the low to mid-range of the autoregulation range. In reality, the vasculature is potentially developed within the low to mid-range of the autoregulation range, consequently tuning its elements to function optimally within this range. As displyed in Fig. 6(a), a larger decrease in blood flow relative to the previous ABNP values is observed in L4 and then L3. Given L4’s initially large flow rate, this reduction brings it closer to the ideal level at physiological ABNP values (50-70 mmHg in our model [86]), while L3 starting near the ideal, shifts further away. Fig. 6(d) demonstrates the mean capillary blood flow at physiological ABNP values, which closely aligns with the data reported by Li et al[9]. This result suggests that the observed reduction in blood flow in layers IV and V of the mouse cerebral vasculature in Li’s study is perhaps caused by the intrinsic limitations in achieving perfect synergy between the vascular segments’ morphological characteristics and the MT mechanism. Furthermore, this analysis demonstrates that blood flow in capillaries across different cortical depths is highly dependent on the ABNP values.

In addition to varying mean capillary blood flow across different cortical depths, simulation results in Fig. 6(a, c) show that the standard deviation of capillary blood flow is larger in superficial layers (L1, L2) compared to deeper layers (L3, L4). This result aligns with the in-vivo observations by Li et al., who reported more uniform capillary blood flow in the deeper cortical layers[9]. In these simulations, our goal was to estimate vessel lumen diameters at various ABNP values to ensure uniform mean blood flow across capillaries. However, there is a systematic distortion in these estimations, which arises from the imperfections in the design of maximally dilated vasculature, estimated RC indices, and the generalization procedure used to model autoregulation. Since IP in superficial segments of PAs is higher than in deeper segments, any distortion in lumen diameters leads to a larger deviation in blood flow in superficial capillaries from the ideal level. As demonstrated in Fig. 6(a), the increase in ABNP and IP in the PA segments of L1 and L2 amplifies these deviations, and more pronounced non-uniformity in blood flow is observed. The same process is potentially happening in reality, with the difference that there is not a single generalized transfer function for the vasculature. Instead, in biological vasculature, each segment, depending on its spatial position within the network, has a unique transfer function that relates the mechanical parameter(s) it detects (input) to an ideal constriction level (output). It is suggested that an increased wall tension (WT) might be directly associated with the potentiation of the myogenic response in blood vessels[92, 93]. This implies that the MT regulatory mechanism in a vascular segment could be calibrated based on the tension exerted on its wall. According to the Laplace’s law, this tension is directly proportional to the IP within that segment, its internal diameter (*D*), and inversely proportional to the segment’s wall thickness (Δ), as expressed in 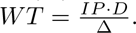 The process of cytoskeletal remodeling or the adjustment of wall thickness in response to wall tension, whether occurring concurrently with SMC contraction to mitigate WT or over extended periods in pathological conditions like hypertension, is not fully understood[92]. Fig. 6(e) displays the relative variation in WT across two segments of a PA located at different cortical depths presuming that vessel wall thickness does not change throughout the autoregulation range. This figure shows that a superficial PA segment experiences a wider range of WT changes compared to a deeper PA. This wide dynamic range of input requires precise transduction into corresponding constriction levels. However, biological systems characterized by high sensitivity, especially across wide ranges of inputs, are subject to larger output distortion. If there is a deviation in how WT is transduced into muscular constriction, whether through the complexity of biological feedback mechanisms[94, 95] or other factors, these deviations are more pronounced in vessels with a wider dynamic range of inputs, such as superficial vessels. Consequently, these larger lumen diameter distortions in superficial layers, when coupled with high IP in these layers, result in a larger mismatch between the ideal balance of IP distribution and the vessel lumen diameters needed to obtain an ideal blood flow regulation. This leads to increased non-uniformity in blood flow in superficial layers, as shown by our simulations and in-vivo experiments[9].

Moreover, Fig. 6(a,c) shows that the standard deviation of blood flow in the capillaries of L4 is larger than in L3. Similarly, measurements by Li et al. of capillary RBC flux across layers I-V in the mouse whisker barrel cortex revealed a larger standard deviation in the capillaries of layer V compared to IV[9]. This large variability in the blood flow of deep capillaries occurs even though the IP is not higher in layer V than in IV. As discussed, high IP not only amplifies the effects of lumen diameter distortion on blood flow variability, but it also means that vessels subjected to high IP are potentially prone to larger deviations in their MT mechanisms, and more distortion in lumen diameter is expected. Therefore, more factors may contribute to the larger variability of blood flow in deep layers. The IP at the entry point of a deep capillary bed relies heavily on the blood flow resistance of PA segments along the penetration direction. According to Poiseuille’s Law, resistance to blood flow in a vessel segment is inversely proportional to the fourth power of its radius, indicating that resistance is highly sensitive to changes in diameter. In deep PA segments, which have smaller diameters, even minor distortions in the lumen diameter result in a relatively large change in resistance. Fig. 6(f) illustrates the mean and standard deviation of blood flow resistance per 10 micrometer length of *PA*_1_ across the four vascular layers at various ABNP values. The data shows that the standard deviation of resistance is larger in L4’s PA segments than in those of the upper layers at various ABNP levels. An explanation for greater variance in resistance of L4’s PA segments lies in the design of our vasculature model. We set the diameter of the deepest PA segment such that it yields an identical WSS to the surface PA segment (Murray’s Law), and then set other PA segments to linearly narrow in between (Sec A2). This simplification fails to yield uniform WSS across all segments, especially considering the specific cortical depths at which capillary beds bifurcate from PAs. This imperfection in the vasculature design adds distortion to the lumen diameters of PA segments. As mentioned, distortion in the lumen diameter of small-diameter vessels, such as deep PA segments, could significantly change vascular segment resistance, and pressure drop across deep PA segments would also be highly sensitive to such distortions. Together, these dynamics would result in a larger standard deviation of capillary blood flow in L4 compared to L3. The amplified impact of lumen diameter distortions on the resistance of small-diameter vascular segments, as observed in our model, can also be applied to deep segments of biological cerebral vasculature, potentially explaining the larger variability observed in deep capillaries. The source of these distortions may arise either from deviations in the MT mechanism or from imperfections in the vasculature design.

In Fig. 5(e,f), the comparison between the ideal and realistic MT profiles indicated that mural cells in mouse cerebral arterioles might not support monophasic flat autoregulation. In the following simulation, we adjusted HRCTF to establish a biphasic ABNP-flow relationship, transitioning from 70 mmHg ABNP to the upper bound of the autoregulation range with a slight slope. The monophasic and biphasic ABNP-flow relationships, along with their corresponding HRCTFs, are displayed in Fig. 6(g). Fig. 6(h) shows the mean and standard deviation of blood flow in non-bifurcating capillaries when vessel lumen diameters were adjusted based on the estimated percent tone for biphasic autoregulation. Although superficial capillaries have slightly higher blood flow at larger ABNP values, in deeper layers blood flow does not decrease below the preferred level, unlike in the monophasic flat autoregulation scenario (Fig. 6(a)). Additionally, Fig. 6(i) displays the relative variations in WT across PA segments located at different cortical depths when the vasculature was modeled for biphasic autoregulation. As shown, WT increases linearly in all segments, unlike the plateaued phase observed at large ABNP values in the middle PA segment under the monophasic flat autoregulation scenario (Fig. 6(e)). This observation suggests that biphasic autoregulation is more logical, since it implies that MT potentiation in all vessels is directly related to the increase in WT. Therefore, the following analyses in this study are based on biphasic autoregulation, consisting of flat phase (ABNP between 40 to 70 mmHg) and sloped phase (ABNP between 70 to 130 mmHg).

Next, we evaluated hemodynamics in various vasculature zones across the entire autoregulation range. Fig. 7(a-c) show the relationships between blood flow, blood flow velocity, and diameter in capillaries. As shown in Fig. 7(a), there is a moderate correlation between blood flow and blood flow velocity, consistent with the theoretical results (Eq. 1) and aligned with in-vivo observations[67, 82, 96, 97]. Fig. 7(b) shows a strong correlation between blood flow and diameter, arising from the adherence to Murray’s law in our vasculature design including in capillaries maximum lumen diameters, and is supported by the experimental data [82]. In contrast, there is no correlation between the blood flow velocity and diameter of capillaries (Fig. 7(c)), which aligns with findings from multiple in-vivo studies [8, 67, 81, 82, 97]. This is while Eq. 1 would suggest a negative correlation, but the incorporation of Murray’s law in our microvasculature design —albeit not in its precise form— offsets this potential negative correlation. These results, in an absence of a noticeable correlation between capillary diameter and blood velocity, suggest that RBC velocity across capillaries is more uniform compared to RBC flux.

**Fig 7.**
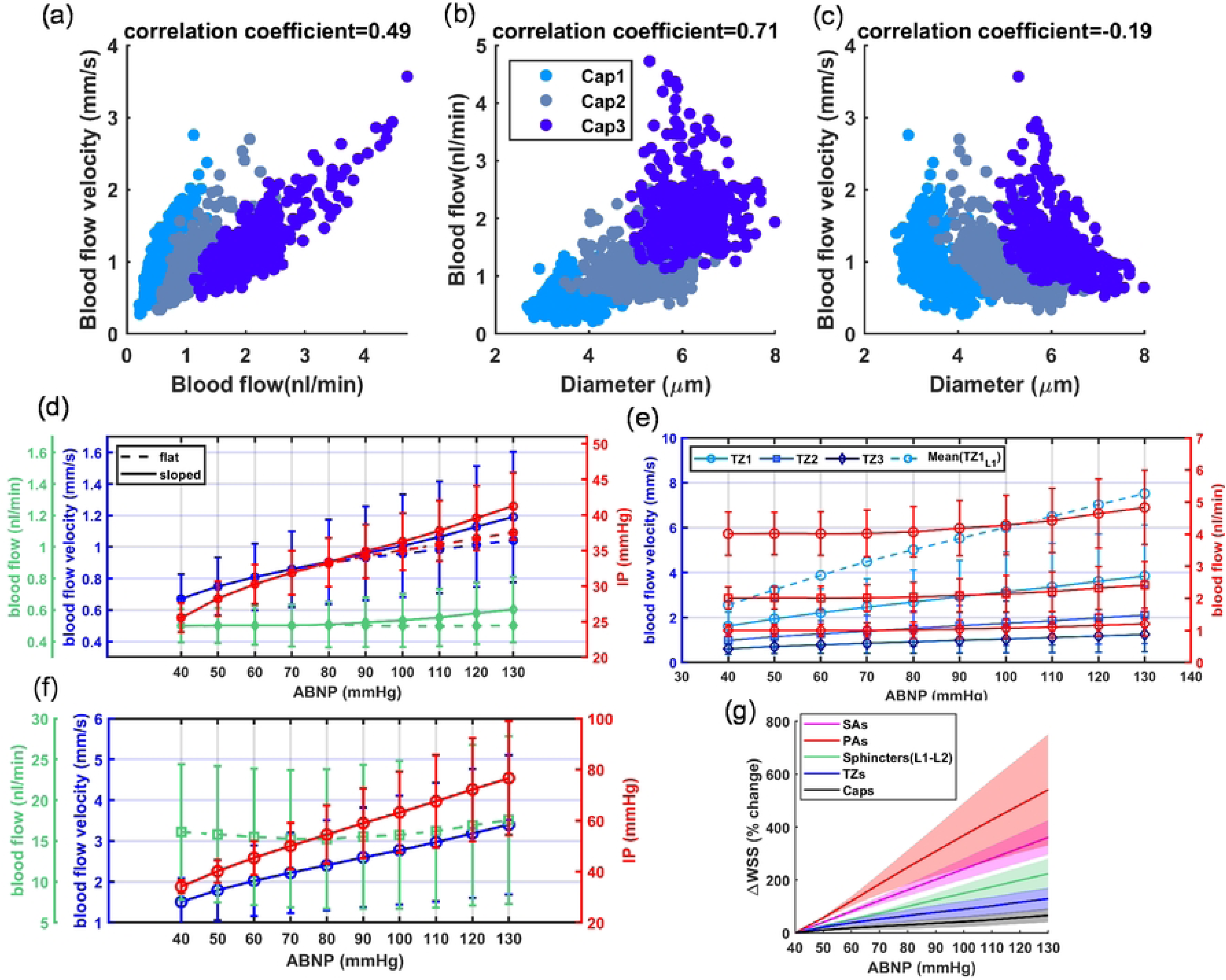
Analysis of hemodynamics across various vasculature zones within the autoregulation range, (a-c) key hemodynamic relationships in capillaries across the autoregulation range: (a) data shows a moderate correlation between blood flow and blood flow velocity, (b) displays a strong correlation between blood flow and diameter, indicating the adherence of our microvasculature design to Murray’s law, (c) shows no correlation between blood flow velocity and luminal diameter in capillaries. Data is collected from a limited number of capillaries across the entire autoregulation range. However, the calculated correlation coefficients are based on the entire database, encompassing 58,800 data points. (d-f) Changes in blood flow, flow velocity, and IP as a function of ABNP for different vascular zones: (d) hemodynamic changes in non-bifurcating capillaries with minimum blood flow levels within the network. The IP axis corresponds to the IP at the entry points of the capillary zone, (e) pronounced variations in blood flow velocity of TZ vessels, which depends on the cortical depth, ABNP, and branching order. Dashed line marks the mean blood flow velocity of TZ1 vessels in L1. Solid lines correspond to similar branching orders across all cortical depths, (f) variability in hemodynamics within PAs across the autoregulation range. The blood supplied to the vasculature, including 30 PAs, is approximately 900 nl/min. This blood supply is nearly thirty times larger than the flow in superficial segments of the PAs and is in close agreement with the data reported by Epp et al. for the similar artery branch[45]. (g) Changes in the WSS across the entire autoregulation range in various vascular zones. The PA zone shows the highest level of WSS increase due to its large contractility and a substantial rise in the apparent viscosity of blood in vessels with PA lumen diameter ranges[71]. Despite nearly uniform WSS in maximally dilated vessels, more pronounced constrictions in segments with larger contractility as ABNP increases have led to an unbalanced increase in WSS in all vessels, challenging the conditions for optimal blood delivery. Solid lines depict the average increase in WSS across various vascular zones, while the shaded areas indicate the standard deviation. The upper boundary of the shaded areas is linked to segments with larger contractility, and the lower boundary is associated with segments with smaller contractility.

Fig. 7(d) illustrates blood flow and blood flow velocity in non-bifurcating capillaries (Cap1), as well as the IP of the entry points of capillary zone (end nodes of TZ segments). The data shows that the mean blood flow in capillaries begins to increase with a slight slope starting at 70 mmHg ABNP (green-solid). The standard deviation of capillary blood flow slightly increases with rising ABNP, mainly due to the larger variability in superficial layers, as shown in Fig. 6(a, c). As these vessels constrict—albeit insignificantly due to their small RC indices—throughout the entire autoregulation range, blood flow velocity does not remain constant and tends to increase with rising ABNP values. This is expected because, according to Eq. 1, a relative decrease in capillary diameter would result in increased velocity. While the reduced level of vessel constriction after 70 mmHg ABNP would lead to a reduced rate of blood flow velocity increase after this point (Fig. 5(e-solid), Fig. 7(d-blue dashed line)), the increase in blood flow in the sloped phase results in a continuous linear increase of blood flow velocity throughout the entire autoregulation range(Fig. 7(d-blue solid line)). This linear dynamic was observed in investigations of cerebral autoregulation in piglet brain pial arterioles[98]. The range of blood flow velocity in this simulation aligns reasonably well with what was calculated in previous computational studies[19, 31, 32] and the RBC speed measured in in-vivo experiments [67, 82, 96, 97]. Notably, in the monophasic flat autoregulation scenario where blood flow remains constant in the network, despite a 225% increase in ABNP values (40-130 mmHg), IP at the capillary entry points does not change as much (Fig. 7(d-red dashed line)), ranging mainly between 25.5 mmHg and 37.5 mmHg (approximately a 47% increase). This relatively small variation in IP at the capillary entry points within the autoregulation range indicates that the IP gradient between the initial nodes of the capillaries and venular nodes, a critical determinant of the force propelling RBCs through capillaries, is primarily regulated by vessels preceding these points. This hemodynamics-vasodynamics interaction in the vasculature underscores the moderate impact of constrictions in capillaries in maintaining constant blood flow within these segments and emphasizes the crucial role of vessels preceding capillaries in accurately regulating IP at these junctures during the autoregulation process. We will delve into a detailed analysis of this aspect of the CBF regulatory mechanism in Sec. B, where we examine distinct contributions of various vasculature zones to the autoregulation.

We proceeded to assess hemodynamics in the TZ vessels across the entire autoregulation range in our vasculature model. Fig. 7(e) shows that blood flow begins to increase with a slight slope starting at 70 mmHg ABNP across all branching orders of the TZ, as expected, though with a larger variability at larger ABNP values. However, blood flow velocity shows noticeable linear changes across the autoregulation range, with the increase being more pronounced in the superficial layers (L1: dashed line in Fig. 7(e), where only the mean values of TZ1 vessels are plotted). The more significant increase in L1 is primarily due to the increased flow in this layer as ABNP rises, shown in Fig. 6(h).This simulation highlights a potential pronounced dependency of RBC speed in TZ vessels on cortical depth and ABNP values; a dynamic yet to be fully explored in in-vivo experiments.

Moreover, we evaluated hemodynamics in other vascular zones. Fig. 7(f) displays the variability of hemodynamics in PAs across the entire autoregulation range. As anticipated, blood flow in PAs also follows the biphasic autoregulation pattern with a large standard deviation. This variability is caused by variations in blood flow at different cortical depths, with superficial segments feeding seven capillary beds having the highest flow and deeper segments the lowest. Similar to TZ vessels, blood flow velocity in PAs is highly dependent on the ABNP value due to the high constriction level in these vessels across the autoregulation range. Additionally, statistical analysis of IP values of PA segments throughout the autoregulation range, as shown in Fig. 7(f), reveals that while the mean and standard deviation of IP increase with rising ABNP, the lower limit of IP (corresponding to deep PA segments) increases at a relatively smaller rate throughout the autoregulation range. This observation suggests that the inherent narrowing of the luminal diameters of PAs toward deeper layers acts similar to sphincters and narrower TZ vessels in superficial layers, serving as a protective mechanism to prevent abnormal high pressure in deep capillaries. This is particularly relevant considering the larger maximum lumen diameter of TZ segments in deeper layers. It can be inferred from this observation that the strategic combination of larger PA segments with narrower TZ segments and sphincters in the superficial layer and their opposite arrangement in deeper layers could help achieve a uniform distribution of capillary blood flow within the layered cerebral vasculature.

Next, we sought to evaluate whether the principle of Murray’s law, which states that optimal blood delivery necessitates a similar level of WSS across vasculature segments, remains valid under conditions where vessels constrict for autoregulation. Although we optimized arteriolar diameters to maintain nearly uniform WSS in arteriolar segments at the maximum dilation state for 40 mmHg ABNP, Fig. 7(g) shows that WSS, calculated based on the simplified Eq. 2, tends to increase more in segments that experience larger constriction (larger RC values). This simulation result suggests that under our systematic evaluation, WSS does not necessarily maintain a uniform level throughout the entire autoregulation range in arteriolar segments, unless there is a regulatory mechanism in the brain vasculature specifically responsible for regulating WSS, a factor not implemented in our analysis. Studies showed that the activation of eNOS by increased WSS could be a key pathway in inhibiting MT and mitigating WSS (refer to Roux et al.[38] study for details). This WSS-dependent inhibition of MT leads to the possibility that the vasculature might employ an additional transfer function, wherein WSS serves as the input and the extent of vessel relaxation as the output. Integrating WSS-dependent inhibition of MT with the WT-dependent potentiation of MT, can theoretically ensure a consistent WSS across all vascular segments, thereby facilitating optimal blood delivery across the autoregulation range. However, such relaxation in vessels, particularly as ABNP increases, could disrupt autoregulation, leading to increased blood flow as constrictions might not be sufficient to maintain nearly constant blood flow. Under these conditions, as ABNP and subsequently blood flow increase, WSS also rises, potentially placing the vasculature in a positive feedback loop at the upper bound of the autoregulation curve. Consequently, the system may non-linearly move toward instability at the upper bound of the autoregulation range, as observed in studies investigating autoregulation in the brain vasculature [1, 98, 99, 100, 101](Fig. 1(c)). This observation suggests that vessels with larger contractility should potentially produce more nitric oxide for vessel relaxation to mitigate the larger increase in WSS throughout the autoregulation range and provide a uniform WSS across all vascular segments. A similar trend was observed recently in the study by Sargent et al., where they reported notably larger endothelial area in cross-sections of TZ vessels with larger contractility compared to the endothelial area in capillaries characterized by smaller contractility [56].

### B: Impact of Different Vascular Zones on CBF Regulatory System: Autoregulation and Functional Hyperemia

In this section, we analyze the lumen diameter-dependent regulation of hemodynamics in the brain vasculature, with emphasis placed on both static (autoregulation) and dynamic (functional hyperemia) conditions. In the previous section, we showed that the product of the vasculature HRCTF and the RC indices of vascular segments provides an estimate of the percent tone in vessels’ segments at ABNP values above 40 mmHg. This estimate reflects the MT level in vessels across the autoregulation range. In this section, our goal is to identify the critical zones in the vasculature where the potentiation or inhibition of MT plays an indispensable role in the optimal regulation of CBF. Fig. 8(a) provides a schematic that illustrates the categorization of vascular zones in this study.

**Fig 8.**
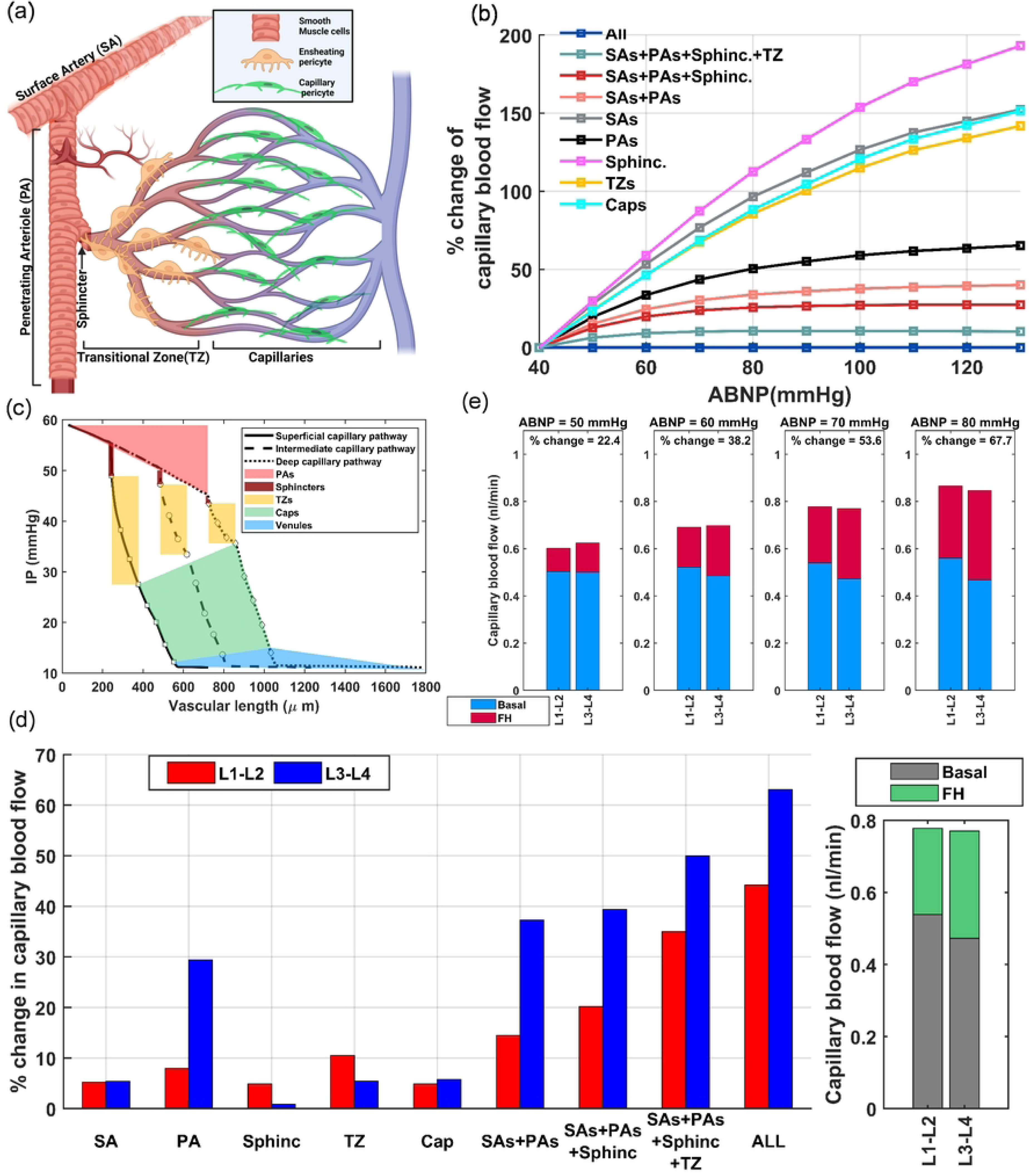
Analyzing the impact of vasodynamics on hemodynamics for CBF regulation across various vascular zones. (a) A schematic representation of various zones within the cerebral vasculature. The figure also illustrates the microvessels’ branching sequence from a PA. This sequence begins with a sphincter, followed by 3-4 layers of diverging branches, each encased by ensheathing pericytes (EPs). The sequence ends in 3-4 layers of converging capillaries, encased by capillary pericytes. Graphics created on BioRender.com, (b) assessing the individual and collective impacts of MT potentiation across various vascular zones on the capillary blood flow. The data shows the relative change in the steady-state mean blood flow in non-bifurcating capillaries under various scenarios, compared to the autoregulated blood flow in the ‘All’ scenario. In the ‘All’ scenario, MT is potentiated in all vascular zones, which results in biphasic autoregulation, (c) pressure drop in three distinct blood flow pathways through superficial, intermediate, and deep capillary networks within the vasculature under 70 mmHg ABNP. The data shows the pressure drop from a surface segment of a PA to a surface segment of an ascending venule. The pressure drop in SAs of our model is about 10 mmHg at 70mmHg ABNP, (d) Left: Distinct and collective impact of MT inhibition across various vascular zones on enhancing blood delivery during FH, assuming NGVC can uniformly inhibit MT in all vascular segments. Right: Mean value of blood flow in superficial and deep capillaries pre-FH and during FH. Although MT inhibition delivers more blood to deep capillaries, since deep layers have lower blood flow pre-FH at 70 mmHg ABNP (Fig 6(h)), blood flow during FH is nearly the same in both deep and superficial layers. (e) Compares the increase in blood flow in response to a 30% MT inhibition in vessels for various ABNP values. Vasculature with a large basal MT shows a more pronounced increase in blood flow in response to a similar MT inhibition. The % change in capillary blood flow for each scenario shows the mean value across both superficial (L1-L2) and deep layers(L3-L4).

#### B1: Autoregulation

Fig. 8(b) shows the relative change in the mean blood flow of non-bifurcating capillaries under various scenarios, compared to autoregulated blood flow when MT is potentiated in all vessels (‘All’ scenario). In this simulation, at each ABNP value, vessel diameters were set based on their estimated percent tone. Depending on the evaluated scenario, we selectively adjusted vessel diameters in specific zones and assessed capillary blood flow. The expectation is that vascular zones with long vascular lengths and large contractility, along with small vessel diameters that generate considerable resistance to increasing flow, would play a key role in the autoregulation. As expected, MT potentiation in PAs alone generally has the largest contribution to autoregulation among other vascular zones. MT potentiation exclusively in SAs, TZs, and Caps can moderate the rise in blood flow almost equally as ABNP increases. Sphincters with their short vascular length contribute the least, primarily in the superficial layers.

According to Poiseuille’s Law, the pressure drop in vascular segments is directly related to the blood flow passing through the segment (Fig. 6(f)). In Fig. 7(d), we showed that, during the autoregulation process, IP at the capillary entry points does not substantially vary throughout the autoregulation range, and IP at these junctures is the primary determinant of blood flow in the vasculature. If IP is increased at these junctures, the blood flow would also increase. However, an increase in blood flow would cause a larger pressure drop in the preceding vascular zones of the capillaries, thereby preventing any further increase in IP at capillary entry points as well as in the flow within the capillaries and the network. IP and flow are coupled systemic parameters due to resistance of the vessels, and the simulation results reflect the steady-state values derived from their balance in the hemodynamic analysis of the network. In Fig. 8(b-PAs), when MT is exclusively potentiated in PAs, blood flow remains relatively constant from the mid-autoregulation range onward. This indicates that the increased blood flow up to the mid-autoregulation range, combined with constriction in PAs, results in a larger pressure drop along the PAs. The augmented IP drop per vessel length under this condition effectively prevents further increases in IP at capillary entry points and limits the increase in the blood flow. This dynamic also explains the declining trend in blood flow increases of all tested scenarios in Fig. 8(b). Therefore, it is impractical to quantify the contribution of various vascular zones to autoregulation. This is because when ABNP increases, and a particular zone either does not contribute or is slow to adjust its myogenic response, blood flow increases beyond the ideal level. In this situation, other zones compensate by generating a higher pressure drop due to the increased blood flow, which helps prevent further increases in blood flow. This process continues until a complete balance between IP, flow, and MT in all vessels is achieved. This dynamic can be observed in experiments performed in Klug’s study, where they abruptly increased the ABNP of mouse retinal vasculature from 20 to 80 mmHg and monitored changes in vessel diameters across different vascular zones [87]. In their experiments, a delayed myogenic response was observed, with the delay being more pronounced in capillaries, followed by TZ vessels, and then arterioles. The variation in the delay of myogenic responses across different vascular zones led to intermittent episodes of constriction and dilation in vessels until a balance between the flow, IP, and MT in all vessels was achieved.

Fig. 8(b) displays the extent of moderating increases in the mean blood flow of non-bifurcating capillaries across all cortical depths if ABNP increases and MT is potentiated in one zone or a combination of zones. However, it does not specifically show the contributions of various vascular zones to the regulation of blood flow at different depths. Fig. 8(c) displays the pressure drop along three distinct blood flow pathways through superficial, intermediate, and deep capillary networks in the vasculature under 70 mmHg ABNP. As shown, there is a pronounced cortical depth-dependency in how different vascular zones regulate blood flow across vascular layers. In superficial layers, sphincters and TZ vessels generate a large pressure drop while PAs generate less. Conversely, in deep layers, PAs produce the largest pressure drop. This suggests that MT in TZ vessels and sphincters plays a more significant role in autoregulating blood flow in superficial layers, while MT in PAs has a more significant role in deep layers.

#### B2: Functional hyperemia

Functional hyperemia refers to the entirety of interactions in brain vasculature that boost blood flow to activated brain regions. This increase in blood flow can be facilitated either by transient vasodilation in the feeding vessels by NGVC or through the deformability of RBCs. Although RBC deformability also plays a role in functional hyperemia [102], our current analysis centers on vasodilation, being the key factor in enhancing capillary blood perfusion. While the biological aspects of NGVC resulting in inhibition of MT in vascular segments are not the focus of this study, our primary goal is to identify vascular zones playing significant roles in augmenting blood flow in capillaries during FH.

In our study of the effects of vasodynamics across various vascular zones on hemodynamics during FH, we began by setting the ABNP to 70 mmHg. At this pressure, vessels have noticeable MT-induced constriction, allowing us to analyze how the inhibition of MT in different vascular zones can facilitate blood delivery. We then assumed that NGVC can inhibit MT by 30% in the vessels. This assumption of uniform inhibition of MT-induced constriction across all vessels is a simplification that does not reflect the realistic physiological conditions of FH for two main reasons. First, different vasodilatory mechanisms operate across various vascular zones [2, 18]. Second, and more importantly, distinct electrophysiological properties of SMCs and pericytes influence the temporal dynamics of MT inhibition by NGVC across different vascular zones [53, 103]. To the best of our knowledge, there is no in-vivo evidence indicating that MT inhibition in capillary pericytes by NGVC occurs swiftly enough for vasodilation to be detected during typical short to moderate FH episodes [57, 104]. These differential temporal dynamics were not considered in our simulation, which analyzes steady-state hemodynamic changes in response to 30% MT inhibition in vessels. In this simulation, we assumed that NGVC inhibits 30% of MT in vessels. However, we estimated the percent tone for vessels at different ABNP values. Therefore, we needed to determine how a 30% inhibition of MT translates to reductions in percent tone and increases in vessel diameters. To identify the relationship between MT and percent tone, we must estimate how vessels passively distend with increases in ABNP under a scenario where mural cells are inactive. In Fig. 5(e), we plotted a passive distension profile of a PA segment at 100 µm cortical depth for various IP values. The curve showing passive distention in relation to IP was remapped to ABNP values and then fitted using an exponential function. Given that this segment is within its maximum active dilation state at an IP range of 27-35 mmHg, we posited that the segment initially reaches this range when ABNP is at 30 mmHg. Note that at this ABNP level, even with vessels at maximum dilation, blood flow falls below the ideal level within the autoregulation range due to the lower IP gradient across the network. From the exponential function fitted to this curve, we derived:

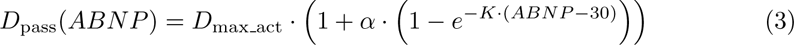

The parameter *α* indicates the percentage increase in passive diameter, *D*_pass_, at high ABNP values (e.g., 130 mmHg) with respect to the maximum active diameter, *D*_max act_. The parameter *K* determines the rate at which the segment passively distends in response to linear increases in ABNP. Through the fitting process, the values of *α* and *K* were determined to be 27.8% and 0.03, respectively. Based on our initial assumption of a similar rate of passive distention in all vessels when ABNP increases from 0 to 40 mmHg until reaching their maximum active dilation state, we extended this assumption to the scenario with inactive mural cells. We assumed vessels continue their passive distention at a similar rate for ABNP values above 40 mmHg, implying a consistent passive distension rate across the network. Therefore, we can generalize the above equation to all vessel segments and write:

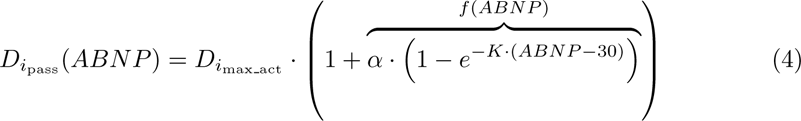

where *i* refers to any segment in the network. Then, by utilizing the definition of percent tone (PT), we can write:

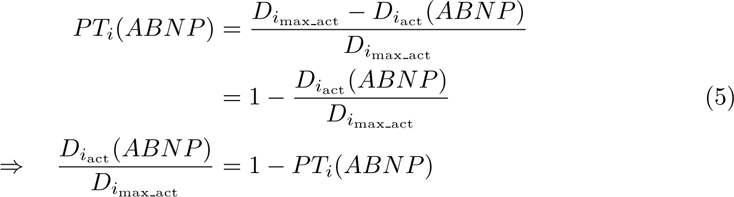

where *D_i_*_max act_ is the active diameter of segment *i* at various ABNP values. Based on the definition of myogenic tone and substituting Eq. 4 and 5 into this definition, the result is:

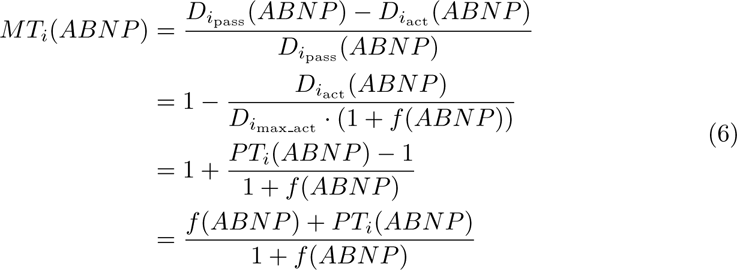

We assumed a 30% inhibition of MT via NGVC:

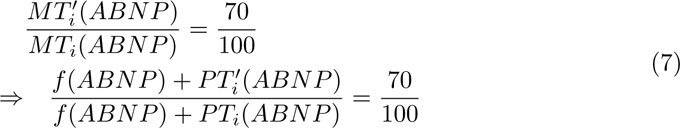

the prime superscript denotes an inhibited or reduced tone, or an increased diameter. By solving the above equation for ABNP=70 and substituting *f* (*ABNP* = 70) = 0.194, we obtain:

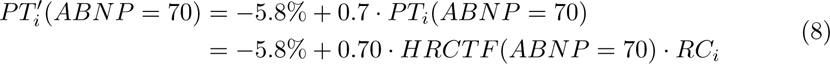

then using Eq. 5 and 8, we can write:

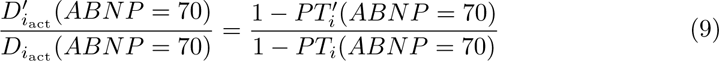

for superficial PA segments with RC indices between 1.0 and 1.1:

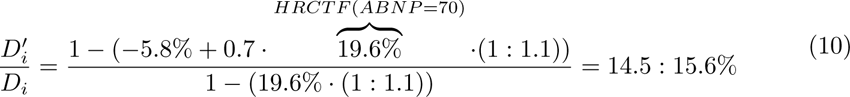

and for capillaries with RC indices around 0.5:

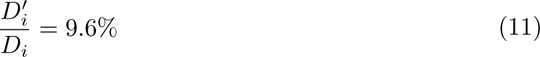

These calculations show that a 30% inhibition of MT does not result in equivalent relative diameter changes across all vessels. For superficial PA segments, it results in approximately a 15% increase in diameter, a range consistent with the relative vasodilation observed in FH experiments in PAs[14, 105, 106, 107].

The results of our simulations for various scenarios are displayed in Fig. 8(d). Considering the differential impact of various vascular zones on hemodynamic regulation across different layers, as shown in Fig. 8(c), we categorized the vasculature into two distinct layers: superficial (L1-L2) and deep (L3-L4). The purpose of this stratification was to identify specific vascular zones that exert a dominant influence on enhancing capillary blood flow in each layer during FH. As expected, the dilation of SAs notably enhances blood flow by lowering the network’s total resistance. Similar to the significant impact of MT potentiation in PAs on moderating blood flow as ABNP increases (Fig. 8(b)), MT inhibition in PAs plays the most substantial role in increasing blood flow in FH, with this effect being more pronounced in deep layers. Conversely, MT inhibition in TZ vessels and sphincters is more impactful on increasing blood flow in superficial layers.

As shown in Fig. 8(c), at physiological levels of ABNP, a significant portion of the pressure drop occurs in capillaries, indicating that they constitute a large portion of the overall vascular resistance in the cerebral vasculature due to their small diameters. This is supported by previous research [19, 34]. However, our simulation results in Fig. 8(d) show that dilation in capillaries minimally enhances blood flow. This suggests that due to the capillaries’ small diameter and the single-file movement of RBCs within them, even an average of 9.6% vasodilation in the capillary zone does not substantially alter the overall resistance of the network to RBCs and blood flow. Additionally, we assumed a 30% MT inhibition in capillaries and an average of 9.6% dilation, which have not been observed during FH in capillaries. This is likely because NGVC-mediated MT inhibition in capillaries either has slow dynamics and cannot achieve a full 30% inhibition during the period of FH, or it only occurs in response to a perceived deficiency of metabolic factors[53]. Assuming a lower percentage of MT inhibition in capillaries, it can be inferred that capillary vasodilation would have an even smaller effect on blood delivery during FH than what is shown in Fig. 8(d). The slow dynamics of MT inhibition in capillaries suggest that to enhance blood flow in capillaries and across the entire vascular system during FH, there are two critical factors: firstly, it involves not the dilation of capillaries but the elevation of IP at their entry points, achieved through the dilation of vessels with high temporal vasoreactivity to NGVC signals; secondly, the reduction of blood viscosity within capillaries, achieved through the deformation of RBCs.

Simulation results in Fig. 8(d) show that vasodilation solely in TZ vessels can increase blood flow by an average of 8%. However, the capacity of TZ vessel dilation to enhance blood flow increases by about 50% when combined with the dilation of preceding vessels. This is evident when comparing blood flow increases in the TZ, SAs+PAs+Sphinc., and SAs+PAs+Sphinc.+TZ scenarios. When TZ vessels undergo dilation, the overall resistance within the network decreases, potentially leading to a proportional increase in the blood flow. However, in the ‘TZ’ scenario, the expected proportional increase in the blood flow is partially counteracted because the blood must traverse through preceding vessels that remain constricted and exert a large resistance. This results in a larger pressure drop before the blood reaches the capillaries and leads to a smaller increase in the steady-state blood flow. These results emphasize that increasing IP at the capillary entry points plays a more determining role in boosting CBF than reducing the overall vascular resistance. Specifically, these simulations show that localized vasodilation, particularly in end-contributor segments such as TZ vessels and capillaries, leads to an increased pressure drop upstream, which can diminish the expected increase in blood flow due to the reduced vascular resistance.

In our next series of simulations, we implemented the same 30% MT inhibition scenario in vessels for various ABNP values. By solving Eq. 7 and calculating the reduced percent tone resulting from the 30% MT inhibition at ABNP=50, 60 and 80 mmHg, we determined the extent of vasodilation in all segments at these ABNP values. Results of these simulations are displayed in Fig. 8(e). Despite the uniform inhibition of MT in all scenarios, the extent of IP drop reduction from the artery boundary node to the capillary entry points differs. For instance, at ABNP of 50 mmHg, MT inhibition led to a 22.4% increase in the average capillary blood perfusion. However, at 80 mmHg, the same inhibition resulted in a 67.7% increase in capillary blood perfusion. These results align with interpretations of the HRCTF profile and the diameter change profile of a PA segment, as shown in Fig. 5(d) and 5(e), respectively. The HRCTF profile shows an initial steep increase at the lower bound of the autoregulation range, with the slope gradually decreasing as the ABNP increases. This pattern indicates that at low ABNP values, vessels must constrict more to achieve the preferred IP drop in vessels leading to capillaries, to ensure preferred capillary blood flow. Conversely, at high ABNP values, even minor vessel constrictions can significantly alter the IP drop. Therefore, MT inhibition at higher ABNP values leads to a more substantial deviation in IP drop in vessels, which significantly increases the blood flow in capillaries.

## Methods

### Mathematical framework for blood flow analysis in the cerebrovascular model

The hemodynamic analysis in this study was based on the research conducted by Secomb’s[72] and Pries’s[51] groups. These teams developed mathematical frameworks that employ empirical laws to enhance the precision and reliability of simulations that model the flow of blood in microvascular networks. These frameworks facilitate the process of assessing blood flow in large cortical networks while incorporating non-Newtonian properties of blood and hematocrit heterogeneity in the model. The mathematical framework was coded in C, and the structural characteristics of the segmented network and hemodynamic parameters were fed as inputs. The flow rate *Q_j_* in segment *j* of the network is presumed to follow Poiseuille’s Law. We had a positive flow in each segment from the start node to the end node. The relationship between nodal pressure *p_k_*and the flow *Q_j_*in segment *j* is formulated by:

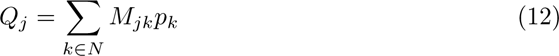

where *N* represents the set of all nodes in the network, and

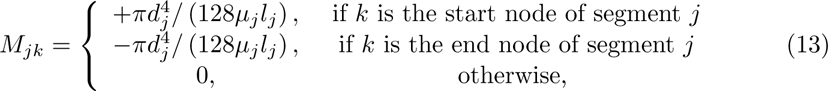

is the conductance between nodes *j* and *k*, with *l_j_, d_j_*, and *µ_j_* signifying the length, diameter, and effective viscosity in segment *j*, respectively. According to mass conservation, the total flow at each interior node is zero. This condition can be combined with the conditions on the boundary nodes to write:

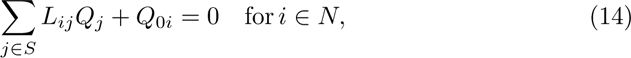

where S denotes all segments in the network, and

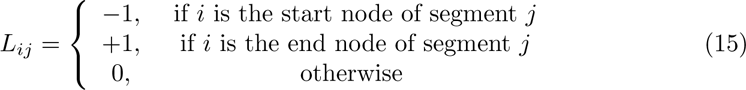

If node *i* is a boundary node, *Q*_0*i*_ indicates the inflow (or outflow if negative). For interior nodes, *Q*_0*i*_ = 0. Combining Eq. 12 and Eq. 14 results in:

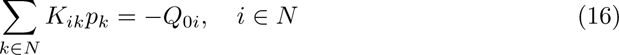

where network conductivity (*K_ik_*) is:

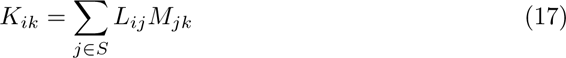

A pressure boundary condition can be enforced at node *i* by substituting the *i*-th row of the matrix *K* with a single diagonal entry of 1 and replacing *Q*_0*i*_ with the specified pressure. If the pressure or flow is known at each boundary node, then the system Eq. 16 is fully determined and can be solved using an iterative process proposed by Secomb et al.[72]. In their proposed mathematical framework of blood flow simulations in microvascular network, the effective viscosity *µ* in segment *j* is a function of the radius *r_j_*and the hematocrit *H_j_*of each segment, utilizing an empirical in-vivo viscosity relationship[108, 109]. The inlet hematocrit was set to 0.4 for pial boundaries. The hematocrit values for all segments can be determined from the flow *Q_j_*, employing empirical relationships for hematocrit partitioning at bifurcation points[109, 110]. To satisfy these relationships, the flow in each vessel was determined using initial values for discharge hematocrit. These flow values were then employed to update *H_j_* in each vessel segment. Effective viscosity was recalculated using the updated hematocrit values. This procedure was repeated until convergence was achieved for *Q_j_*, *H_j_*, and *µ_j_* in each segment.

### Statistical Analysis

In Sec. A1, we performed statistical analysis on two-photon images of murine cerebral vasculature to investigate the location of sphincters and estimate the diameters of daughter and parent vessels at sphincter sites. In the algorithm we employed, we traced arterioles’ path as they penetrate the brain and marked bifurcation nodes. We then searched for patterns resembling sphincter signatures—constriction succeeded by bulb-shaped distension—to confirm the presence of a sphincter. Following successful detection, we estimated the diameter ratio of the daughter to parent branch. For the diameter quantification of TZ vessels, our analysis relied on existing measurements and branching order data provided by the database from Anna Devor’s laboratory[47, 48]. The findings presented in Fig 2(c) are entirely based on this external dataset, which enabled us to incorporate more detailed morphological characteristics in our model.

## Discussion

In this work, we present a computational investigation of hemodynamic-vasodynamic interactions within cerebral vasculature to model autoregulation. Blood vessel segments, due to their myogenic response regulatory mechanisms, function as variable hydraulic resistors. These vessel segments adjust their diameters autonomously in response to mechanical forces exerted by the flowing blood to maintain constant flow in capillaries. In our study, we employed various simplifications and generalizations to capture these dynamics. First, we assumed that as the mean arterial pressure increases from zero, the vasculature begins to inflate due to passive distention in the vessels. Next, we assumed that the maximum dilation state for all vessels occurs when the ABNP in the model reaches 40 mmHg, which is a hypothetical value that falls within the lower limit of the autoregulation range. The analysis of the model shows that to maintain the uniformity in capillary blood flow at various cortical depths in the maximally dilated state, a set of morphological features need to be incorporated in the model’s design. For example, sphincters and narrower TZ segments should be included in the structure of the upper microvascular layers. Furthermore, a progressive narrowing of arteries and arterioles, as these vessels bifurcate from the main brain feeding arteries and penetrate deeper into the tissue, also contributes to the uniformity of capillary blood flow. When the ABNP exceeds 40 mmHg, the vascular-centric myogenic response helps the network to obtain uniform blood flow in capillaries by adjusting the vessel constriction levels. In our model, we introduced the relative contractility index to quantify the expression level of contractile elements in each vascular segment. We assigned this index to all segments and showed that elevating ABNP from 40 mmHg to 130 mmHg, when paired with a well-regulated myogenic response in all vessels, can effectively model blood flow autoregulation across all cortical depths (Fig. 6(a,h)).

We investigated the validity of our computational model by comparing its dynamics to experimental observations. Consistent with experimental data[9], our model showed that the blood flow in middle-layer capillaries is less than the flow in both deep and superficial layers at physiological ABNP values (Fig. 6(a,d,h)). This variability of blood flow as a function of cortical depth arises from the complexity in interactions among the morphological characteristics and the adjustments of MT in all vascular segments within the broad autoregulation range. We showed that only in a narrow band of the autoregulation range it is possible to achieve a uniform mean blood flow in capillaries at all depths. Furthermore, our model showed that superficial and deep capillaries tend to have more non-uniform blood flow compared to middle-layer capillaries (Fig. 6(c)), a finding that is consistent with in-vivo observations [9]. Moreover, our analysis revealed that blood flow velocity in capillaries remains relatively stable, contrasting with the pronounced variability observed in TZ vessels[8, 104], sphincters, and PAs. We showed that this variability is dependent on cortical depth and the vasculature’s position within its autoregulation range.

After modeling and validating the autoregulation in our cerebrovascular model, we shifted our focus to analyze hemodynamics across different vascular zones throughout the autoregulation range(Fig. 7). First, we identified that IP at the entry points of the capillary bed is a critical factor of the autoregulation (Fig. 7(d)). We found that the precise development of myogenic tone in vessels leading to these entry points is crucial for regulating this key variable. Our analysis revealed the dependency on cortical depth in this regulation: in superficial layers, TZ vessels and sphincters play a major role, with a lesser contribution from PAs. In contrast, this contribution pattern is reversed in deeper layers. This depth-dependent dynamic was clearly evident in our simulation results (Fig. 8(c)).

Similarly, the depth-dependent contribution of various vascular zones through MT potentiation for autoregulation can be extended to MT inhibition by NGVC during FH to enhance blood delivery. In superficial layers, MT inhibition (vasodilation) in sphincters and TZ vessels leads to a larger increase in blood flow, while MT inhibition in PAs substantially increases blood flow in deep capillaries (Fig. 8(d)). However, unlike the static nature of autoregulation, FH is a dynamic process, meaning we cannot simply quantify the contribution of various vascular zones to blood delivery during FH without considering factors such as the intensity and duration of neuronal activity, pre-FH MT levels in vessels, or distinct differences in mural cell biology across various vascular zones that can affect the extent and timing of MT inhibition by NGVC. For example, in Fig. 8(e), we showed that equal levels of MT inhibition via NGVC at higher pre-FH MT levels led to a larger increase in blood flow; however, if all environmental factors in FH experiments remain constant, except for the IP in the main brain-feeding arteries, and similar neuronal activity is induced, it is uncertain whether NGVC would inhibit MT to the same extent at different pre-FH MT levels. Mural cell biology suggests that they may adapt to manage these variations and prevent excessive or insufficient blood delivery during FH, regardless of pre-FH MT levels. It is now well-recognized that MT-induced constriction in mural cells is primarily controlled by WT-dependent depolarizing ion channels[92]. On the other hand, rapid inhibition of MT during FH is likely to be achieved through the direct induction of hyperpolarization in mural cells caused by adjacent cells such as endothelial cells [111, 112], astrocytes [105, 113], and neurons[114] and not by suppressing the activity of depolarizing ion channels of mural cells. The degree of MT inhibition through induced hyperpolarization in mural cells by adjacent cells depends on the ability of the hyperpolarizing currents to counteract WT-depolarizing currents. When WT is large, depolarizing channels are more active, the resting membrane resistance of mural cells is smaller, and hyperpolarizing factors have a reduced capacity to hyperpolarize mural cells and inhibit MT. Given these uncertainties regarding the timing and extent of MT inhibition via NGVC during FH in different vascular zones, it is technically challenging to quantify the contribution of these zones to FH. Although our model did not explicitly address this dynamic, we can infer that regions with small-diameter vessels, high contractility, and strong vasoreactivity to NGVC—such as PAs, sphincters, and TZs—are likely to contribute more than other vascular zones.

In a healthy brain, regardless of the level of neuronal activity, MT development in vascular segments will bias the network to an appropriate operating point for the RBC flux through capillaries (0.5 nl/min in Cap1 in our model). Our computational analysis supports quadriphasic ABNP-flow dynamics that have been previously observed experimentally[98], including the following physiological phases: low ABNP with flow below the operating point, moderate ABNP with flow near the operating point (flat phase of autoregulation), high ABNP with controlled increases at the operating point (sloped phase of autoregulation), and extremely high ABNP with uncontrolled flow increases (unstable region). These phases are regulated by vascular cells to mitigate hemodynamic forces, such as WT and WSS, which could otherwise compromise vessel wall integrity. We provided computational evidence supporting that an increase in WT caused by increments in mean arteriolar pressure can be mitigated by MT in vessels and our model suggests that this property is linear. Furthermore, a reduction in capacity of mural cells to induce a large constriction in the mid to upper range of the autoregulation (attributed to endothelial cells’ efforts to mitigate increased WSS) prevents the formation of a plateau throughout the autoregulatory range. Focusing within the autoregulation range, our computational analysis indicates that the biphasic ABNP-blood flow dynamic is the result of the synergy between the morphological (e.g., vessel lumen diameter and expression levels of contractile elements) and the mechanobiological (MT adjustment by vascular cells and regulated by hemodynamics) characteristics of the vascular network. This biphasic dynamic in the ABNP-flow results in a monophasic, linear ABNP-blood flow velocity dynamic across vessels throughout the autoregulation range, again aligning with experimental observations[98, 115].

Our coarsely segmented computational model of the cerebrovasculature required that we make several simplifications to streamline cerebral autoregulation modeling that may have had a bearing on outcomes. First, the capillary network within our microvessel structure was modeled by a simplified hierarchy of 2-to-1 converging vessels. Although this model can emulate reasonably accurate flow and pressure within the network, it does not capture the net-like structure of capillaries in their full complexity [116, 117]. In particular, the three distinct decremental trends observed in the capillary blood flow velocity-diameter relationship (Fig. 4(c)) are likely a result of this simplification. While our capillary network is not a net-like model, when we incorporated Murray’s Law in the microvasculature design, no pronounced correlation in the capillary blood flow velocity-diameter was observed, similar to what we see in reality[8, 67, 81, 82, 97]. This means that the three distinct decremental trends in our simplified model (Fig. 4(c)) do not fully replicate the hemodynamics in a realistic net-like structure of capillaries. Second, our approach in assigning relative contractility to each segment of our cerebrovasculature model was based on a plausible yet speculative assumption: segments subject to higher IP in maximally dilated vasculature were presumed to be encased by a denser array of contractile elements. The advent of advanced imaging technologies now offers the potential to quantify different morphological characteristics of vascular segments, such as luminal diameter, vessel wall thickness, and the expression level of contractile elements, more accurately and realistically, promising improved accuracy in future models.

In summary, the proposed simplified model provides a computationally effective framework for multi-scale studies. It features a closed, coarsely segmented, circulatory network and integrates hemodynamics (WT and WSS) as fundamental inputs in the transfer functions of vasodynamics. This design makes it possible to model dynamic adjustments in vessel lumen diameters including both delayed and active forces generated by muscle activity and the immediate passive distension. Our computational framework can be expanded to more sophisticated multi-scale computational studies, where broader network inputs (e.g., neuron or astrocyte derived signaling) might be incorporated to model, interrogate, and analyze transient hemodynamic/vasodynamic changes in the cerebrovasculature.

## Acknowledgments

This work was supported by NIH/NIA grant R01AG067330, the National Science Foundation (NSF) grants 2154267, 1830145, and by the Army Research Office (ARO) grant W911NF1810323. This study was conducted using resources and facilities at the William S. Middleton Memorial Veterans Hospital, Madison, WI.

## References

1. Claassen JA, Thijssen DH, Panerai RB, Faraci FM. Regulation of cerebral blood flow in humans: physiology and clinical implications of autoregulation. Physiological reviews. 2021;101(4):1487–1559.

2. Iadecola C. The Neurovascular Unit Coming of Age: A Journey through Neurovascular Coupling in Health and Disease. Neuron. 2017;96(1):17–42. 10.1016/j.neuron.2017.07.030.

3. Nortley R, Korte N, Izquierdo P, Hirunpattarasilp C, Mishra A, Jaunmuktane Z, et al. Amyloid *β* oligomers constrict human capillaries in Alzheimer’s disease via signaling to pericytes. Science. 2019;365(6450):eaav9518.

4. Korte N, Ilkan Z, Pearson CL, Pfeiffer T, Singhal P, Rock JR, et al. The Ca 2+-gated channel TMEM16A amplifies capillary pericyte contraction and reduces cerebral blood flow after ischemia. The Journal of Clinical Investigation. 2022;132(9).

5. Stobart JL, Erlebach E, Glück C, Huang SF, Barrett MJ, Li M, et al. Altered hemodynamics and vascular reactivity in a mouse model with severe pericyte deficiency. Journal of Cerebral Blood Flow & Metabolism. 2023;43(5):763–777.

6. Munting LP, Derieppe M, Suidgeest E, Hirschler L, Van Osch MJ, Denis de Senneville B, et al. Cerebral blood flow and cerebrovascular reactivity are preserved in a mouse model of cerebral microvascular amyloidosis. Elife. 2021;10:e61279.

7. Fang X, Tang C, Zhang H, Border JJ, Liu Y, Shin SM, et al. Longitudinal characterization of cerebral hemodynamics in the TgF344-AD rat model of Alzheimer’s disease. GeroScience. 2023; p. 1–20.

8. Watson AN, Berthiaume AA, Faino AV, McDowell KP, Bhat NR, Hartmann DA, et al. Mild pericyte deficiency is associated with aberrant brain microvascular flow in aged PDGFR*β*+/- mice. Journal of Cerebral Blood Flow & Metabolism. 2020;40(12):2387–2400.

9. Li B, Esipova TV, Sencan I, Kılıç K, Fu B, Desjardins M, et al. More homogeneous capillary flow and oxygenation in deeper cortical layers correlate with increased oxygen extraction. Elife. 2019;8:e42299.

10. Marchand PJ, Lu X, Zhang C, Lesage F. Validation of red blood cell flux and velocity estimations based on optical coherence tomography intensity fluctuations. Scientific Reports. 2020;10(1):19584.

11. Li B, Ohtomo R, Thunemann M, Adams SR, Yang J, Fu B, et al. Two-photon microscopic imaging of capillary red blood cell flux in mouse brain reveals vulnerability of cerebral white matter to hypoperfusion. Journal of Cerebral Blood Flow & Metabolism. 2020;40(3):501–512.

12. Berthiaume AA, Schmid F, Stamenkovic S, Coelho-Santos V, Nielson CD, Weber B, et al. Pericyte remodeling is deficient in the aged brain and contributes to impaired capillary flow and structure. Nature Communications. 2022;13(1):5912.

13. Kurtz TW, Lujan HL, DiCarlo SE. The 24 h pattern of arterial pressure in mice is determined mainly by heart rate-driven variation in cardiac output. Physiological Reports. 2014;2(11):e12223.

14. Park L, Hochrainer K, Hattori Y, Ahn SJ, Anfray A, Wang G, et al. Tau induces PSD95–neuronal NOS uncoupling and neurovascular dysfunction independent of neurodegeneration. Nature neuroscience. 2020;23(9):1079–1089.

15. Van Skike CE, Hussong SA, Hernandez SF, Banh AQ, DeRosa N, Galvan V. mTOR Attenuation with Rapamycin Reverses Neurovascular Uncoupling and Memory Deficits in Mice Modeling Alzheimer’s Disease. Journal of Neuroscience. 2021;41(19):4305–4320.

16. Royea J, Martinot P, Hamel E. Memory and cerebrovascular deficits recovered following angiotensin IV intervention in a mouse model of Alzheimer’s disease. Neurobiology of Disease. 2020;134:104644.

17. Shabir O, Sharp P, Rebollar MA, Boorman L, Howarth C, Wharton SB, et al. Enhanced cerebral blood volume under normobaric hyperoxia in the J20-hAPP mouse model of Alzheimer’s disease. Scientific reports. 2020;10(1):1–10.

18. Pfeiffer T, Li Y, Attwell D. Diverse mechanisms regulating brain energy supply at the capillary level. Current Opinion in Neurobiology. 2021;69:41–50.

19. Gould IG, Tsai P, Kleinfeld D, Linninger A. The capillary bed offers the largest hemodynamic resistance to the cortical blood supply. Journal of Cerebral Blood Flow & Metabolism. 2017;37(1):52–68.

20. Hall CN, Reynell C, Gesslein B, Hamilton NB, Mishra A, Sutherland BA, et al. Capillary pericytes regulate cerebral blood flow in health and disease. Nature. 2014;508(7494):55–60.

21. Rungta RL, Chaigneau E, Osmanski BF, Charpak S. Vascular compartmentalization of functional hyperemia from the synapse to the pia. Neuron. 2018;99(2):362–375.

22. Lampe R, Botkin N, Turova V, Blumenstein T, Alves-Pinto A, et al. Mathematical modelling of cerebral blood circulation and cerebral autoregulation: towards preventing intracranial hemorrhages in preterm newborns. Computational and mathematical methods in medicine. 2014;2014.

23. Carlson BE, Arciero JC, Secomb TW. Theoretical model of blood flow autoregulation: roles of myogenic, shear-dependent, and metabolic responses. American Journal of Physiology-Heart and Circulatory Physiology. 2008;295(4):H1572–H1579.

24. Piechnik SK, Chiarelli PA, Jezzard P. Modelling vascular reactivity to investigate the basis of the relationship between cerebral blood volume and flow under CO2 manipulation. Neuroimage. 2008;39(1):107–118.

25. Ursino M, Lodi CA. A simple mathematical model of the interaction between intracranial pressure and cerebral hemodynamics. Journal of applied physiology. 1997;82(4):1256–1269.

26. Banaji M, Tachtsidis I, Delpy D, Baigent S. A physiological model of cerebral blood flow control. Mathematical biosciences. 2005;194(2):125–173.

27. Daher A, Payne S. A network-based model of dynamic cerebral autoregulation. Microvascular Research. 2023;147:104503.

28. Ferrandez A, David T, Brown M. Numerical models of auto-regulation and blood flow in the cerebral circulation. Computer Methods in Biomechanics & Biomedical Engineering. 2002;5(1):7–19.

29. Spronck B, Martens EG, Gommer ED, van de Vosse FN. A lumped parameter model of cerebral blood flow control combining cerebral autoregulation and neurovascular coupling. American Journal of Physiology-Heart and Circulatory Physiology. 2012;303(9):H1143–H1153.

30. Ii S, Kitade H, Ishida S, Imai Y, Watanabe Y, Wada S. Multiscale modeling of human cerebrovasculature: A hybrid approach using image-based geometry and a mathematical algorithm. PLoS computational biology. 2020;16(6):e1007943.

31. Sweeney PW, Walker-Samuel S, Shipley RJ. Insights into cerebral haemodynamics and oxygenation utilising in vivo mural cell imaging and mathematical modelling. Scientific Reports. 2018;8(1):1–15.

32. Gagnon L, Smith AF, Boas DA, Devor A, Secomb TW, Sakadžić S. Modeling of cerebral oxygen transport based on in vivo microscopic imaging of microvascular network structure, blood flow, and oxygenation. Frontiers in computational neuroscience. 2016;10:82.

33. Hartung G, Vesel C, Morley R, Alaraj A, Sled J, Kleinfeld D, et al. Simulations of blood as a suspension predicts a depth dependent hematocrit in the circulation throughout the cerebral cortex. PLoS computational biology. 2018;14(11):e1006549.

34. Schmid F, Tsai PS, Kleinfeld D, Jenny P, Weber B. Depth-dependent flow and pressure characteristics in cortical microvascular networks. PLoS computational biology. 2017;13(2):e1005392.

35. Linninger A, Hartung G, Badr S, Morley R. Mathematical synthesis of the cortical circulation for the whole mouse brain-part I. theory and image integration. Computers in biology and medicine. 2019;110:265–275.

36. Reichold J, Stampanoni M, Keller AL, Buck A, Jenny P, Weber B. Vascular graph model to simulate the cerebral blood flow in realistic vascular networks. Journal of Cerebral Blood Flow & Metabolism. 2009;29(8):1429–1443.

37. Fang Q, Sakadžić S, Ruvinskaya L, Devor A, Dale AM, Boas DA. Oxygen advection and diffusion in a three-dimensional vascular anatomical network. Optics express. 2008;16(22):17530–17541.

38. Roux E, Bougaran P, Dufourcq P, Couffinhal T. Fluid shear stress sensing by the endothelial layer. Frontiers in Physiology. 2020;11:861.

39. Humphrey JD, Schwartz MA. Vascular mechanobiology: homeostasis, adaptation, and disease. Annual review of biomedical engineering. 2021;23:1–27.

40. Taber LA. An optimization principle for vascular radius including the effects of smooth muscle tone. Biophysical journal. 1998;74(1):109–114.

41. Coelho-Santos V, Shih AY. Postnatal development of cerebrovascular structure and the neurogliovascular unit. Wiley Interdisciplinary Reviews: Developmental Biology. 2020;9(2):e363.

42. Novák B, Tyson JJ. Design principles of biochemical oscillators. Nature reviews Molecular cell biology. 2008;9(12):981–991.

43. Kenny A, Zakkaroff C, Plank MJ, David T. Massively parallel simulations of neurovascular coupling with extracellular diffusion. Journal of computational science. 2018;24:116–124.

44. Blinder P, Tsai PS, Kaufhold JP, Knutsen PM, Suhl H, Kleinfeld D. The cortical angiome: an interconnected vascular network with noncolumnar patterns of blood flow. Nature neuroscience. 2013;16(7):889–897.

45. Epp R, Glück C, Binder NF, El Amki M, Weber B, Wegener S, et al. The role of leptomeningeal collaterals in redistributing blood flow during stroke. PLoS computational biology. 2023;19(10):e1011496.

46. Adams MD, Winder AT, Blinder P, Drew PJ. The pial vasculature of the mouse develops according to a sensory-independent program. Scientific reports. 2018;8(1):9860.

47. Uhlirova H, Tian P, Kılıç K, Thunemann M, Sridhar VB, Bartsch H, et al. Neurovascular Network Explorer 2.0: a database of 2-photon single-vessel diameter measurements from mouse SI cortex in response to optogenetic stimulation. Frontiers in neuroinformatics. 2017; p. 4.

48. Sridhar VB, Tian P, Dale AM, Devor A, Saisan PA. Neurovascular Network Explorer 1.0: a database of 2-photon single-vessel diameter measurements with MATLAB® graphical user interface. Frontiers in neuroinformatics. 2014;8:56.

49. Zambach SA, Cai C, Helms HCC, Hald BO, Dong Y, Fordsmann JC, et al. Precapillary sphincters and pericytes at first-order capillaries as key regulators for brain capillary perfusion. Proceedings of the National Academy of Sciences. 2021;118(26).

50. Grubb S, Lauritzen M, Aalkjær C. Brain capillary pericytes and neurovascular coupling. Comparative Biochemistry and Physiology Part A: Molecular & Integrative Physiology. 2021;254:110893.

51. Pries AR, Secomb TW. Blood flow in microvascular networks. In: Microcirculation. Elsevier; 2008. p. 3–36.

52. Ji X, Ferreira T, Friedman B, Liu R, Liechty H, Bas E, et al. Brain microvasculature has a common topology with local differences in geometry that match metabolic load. Neuron. 2021;109(7):1168–1187.

53. Hartmann DA, Berthiaume AA, Grant RI, Harrill SA, Koski T, Tieu T, et al. Brain capillary pericytes exert a substantial but slow influence on blood flow. Nature neuroscience. 2021;24(5):633–645.

54. Longden TA, Mughal A, Hennig GW, Harraz OF, Shui B, Lee FK, et al. Local IP3 receptor–mediated Ca2+ signals compound to direct blood flow in brain capillaries. Science Advances. 2021;7(30):eabh0101.

55. Ornelas S, Berthiaume AA, Bonney SK, Coelho-Santos V, Underly RG, Kremer A, et al. Three-dimensional ultrastructure of the brain pericyte-endothelial interface. Journal of Cerebral Blood Flow & Metabolism. 2021;41(9):2185–2200.

56. Sargent SM, Bonney SK, Li Y, Stamenkovic S, Takeno MM, Coelho-Santos V, et al. Endothelial structure contributes to heterogeneity in brain capillary diameter. Vascular Biology. 2023;5(1).

57. Suzuki H, Takeda H, Takuwa H, Ji B, Higuchi M, Kanno I, et al. Capillary responses to functional and pathological activations rely on the capillary states at rest. Journal of Cerebral Blood Flow & Metabolism. 2023;43(6):1010–1024.

58. Grant RI, Hartmann DA, Underly RG, Berthiaume AA, Bhat NR, Shih AY. Organizational hierarchy and structural diversity of microvascular pericytes in adult mouse cortex. Journal of Cerebral Blood Flow & Metabolism. 2019;39(3):411–425.

59. Gutiérrez-Jiménez E, Cai C, Mikkelsen IK, Rasmussen PM, Angleys H, Merrild M, et al. Effect of electrical forepaw stimulation on capillary transit-time heterogeneity (CTH). Journal of Cerebral Blood Flow & Metabolism. 2016;36(12):2072–2086.

60. Cai C, Fordsmann JC, Jensen SH, Gesslein B, Lønstrup M, Hald BO, et al. Stimulation-induced increases in cerebral blood flow and local capillary vasoconstriction depend on conducted vascular responses. Proceedings of the National Academy of Sciences. 2018;115(25):E5796–E5804.

61. Steinman J, Koletar MM, Stefanovic B, Sled JG. 3D morphological analysis of the mouse cerebral vasculature: Comparison of in vivo and ex vivo methods. PloS one. 2017;12(10):e0186676.

62. Mughal A, Nelson MT, Hill-Eubanks D. The post-arteriole transitional zone: a specialized capillary region that regulates blood flow within the CNS microvasculature. The Journal of Physiology. 2023;601(5):889–901.

63. Ghanavati S, Lerch JP, Sled JG. Automatic anatomical labeling of the complete cerebral vasculature in mouse models. Neuroimage. 2014;95:117–128.

64. Xiong B, Li A, Lou Y, Chen S, Long B, Peng J, et al. Precise cerebral vascular atlas in stereotaxic coordinates of whole mouse brain. Frontiers in neuroanatomy. 2017;11:128.

65. Meng G, Zhong J, Zhang Q, Wong JS, Wu J, Tsia KK, et al. Ultrafast two-photon fluorescence imaging of cerebral blood circulation in the mouse brain in vivo. Proceedings of the National Academy of Sciences. 2022;119(23):e2117346119.

66. Lyons DG, Parpaleix A, Roche M, Charpak S. Mapping oxygen concentration in the awake mouse brain. Elife. 2016;5:e12024.

67. Kleinfeld D, Mitra PP, Helmchen F, Denk W. Fluctuations and stimulus-induced changes in blood flow observed in individual capillaries in layers 2 through 4 of rat neocortex. Proceedings of the National Academy of Sciences. 1998;95(26):15741–15746.

68. Wälchli T, Bisschop J, Miettinen A, Ulmann-Schuler A, Hintermüller C, Meyer EP, et al. Hierarchical imaging and computational analysis of three-dimensional vascular network architecture in the entire postnatal and adult mouse brain. Nature protocols. 2021;16(10):4564–4610.

69. Bennett RE, Robbins AB, Hu M, Cao X, Betensky RA, Clark T, et al. Tau induces blood vessel abnormalities and angiogenesis-related gene expression in P301L transgenic mice and human Alzheimer’s disease. Proceedings of the National Academy of Sciences. 2018;115(6):E1289–E1298.

70. Murray CD. The physiological principle of minimum work: I. The vascular system and the cost of blood volume. Proceedings of the National Academy of Sciences. 1926;12(3):207–214.

71. Secomb TW, Pries AR. Blood viscosity in microvessels: experiment and theory. Comptes Rendus Physique. 2013;14(6):470–478.

72. Fry BC, Lee J, Smith NP, Secomb TW. Estimation of blood flow rates in large microvascular networks. Microcirculation. 2012;19(6):530–538.

73. Brunner C, Macé E, Montaldo G, Urban A. Quantitative hemodynamic measurements in cortical vessels using functional ultrasound imaging. Frontiers in Neuroscience. 2022;16:831650.

74. Coelho-Santos V, Berthiaume AA, Ornelas S, Stuhlmann H, Shih AY. Imaging the construction of capillary networks in the neonatal mouse brain. Proceedings of the National Academy of Sciences. 2021;118(26):e2100866118.

75. Nguyen J, Nishimura N, Fetcho RN, Iadecola C, Schaffer CB. Occlusion of cortical ascending venules causes blood flow decreases, reversals in flow direction, and vessel dilation in upstream capillaries. Journal of Cerebral Blood Flow & Metabolism. 2011;31(11):2243–2254.

76. Qi Y, Roper M. Control of low flow regions in the cortical vasculature determines optimal arterio-venous ratios. Proceedings of the National Academy of Sciences. 2021;118(34):e2021840118.

77. Bonney S, Sullivan L, Cherry T, Daneman R, Shih A. Distinct features of brain perivascular fibroblasts and mural cells revealed by in vivo two-photon imaging. bioRxiv. Preprint]. 2021;10(2021.05):14–444194.

78. Qiu B, Zhao Z, Wang N, Feng Z, Chen Xj, Chen W, et al. A systematic observation of vasodynamics from different segments along the cerebral vasculature in the penumbra zone of awake mice following cerebral ischemia and recanalization. Journal of Cerebral Blood Flow & Metabolism. 2023;43(5):665–679.

79. Taylor ZJ, Hui ES, Watson AN, Nie X, Deardorff RL, Jensen JH, et al. Microvascular basis for growth of small infarcts following occlusion of single penetrating arterioles in mouse cortex. Journal of Cerebral Blood Flow & Metabolism. 2016;36(8):1357–1373.

80. Kucharz K, Kristensen K, Johnsen KB, Lund MA, Lønstrup M, Moos T, et al. Post-capillary venules are the key locus for transcytosis-mediated brain delivery of therapeutic nanoparticles. Nature communications. 2021;12(1):4121.

81. Li B, Lu X, Moeini M, Sakadžić S, Thorin E, Lesage F. Atherosclerosis is associated with a decrease in cerebral microvascular blood flow and tissue oxygenation. PLoS One. 2019;14(8):e0221547.

82. Desjardins M, Berti R, Lefebvre J, Dubeau S, Lesage F. Aging-related differences in cerebral capillary blood flow in anesthetized rats. Neurobiology of aging. 2014;35(8):1947–1955.

83. Santisakultarm TP, Cornelius NR, Nishimura N, Schafer AI, Silver RT, Doerschuk PC, et al. In vivo two-photon excited fluorescence microscopy reveals cardiac-and respiration-dependent pulsatile blood flow in cortical blood vessels in mice. American Journal of Physiology-Heart and Circulatory Physiology. 2012;302(7):H1367–H1377.

84. Hartmann DA, Coelho-Santos V, Shih AY. Pericyte Control of Blood Flow Across Microvascular Zones in the Central Nervous System. Annual Review of Physiology. 2022;84(1):331–354. doi:10.1146/annurev-physiol-061121-040127.

85. Grubb S. Ultrastructure of precapillary sphincters and the neurovascular unit. Vascular Biology. 2023;5(1).

86. Jeffrey DA, Russell A, Bueno Guerrero M, Fontaine JT, Romero P, Rosehart AC, et al. Estrogen regulates myogenic tone in hippocampal arterioles by enhanced basal release of nitric oxide and endothelial SKCa channel activity. bioRxiv. 2023; p. 2023–08.

87. Klug NR, Sancho M, Gonzales AL, Heppner TJ, O’Brien RIC, Hill-Eubanks D, et al. Intraluminal pressure elevates intracellular calcium and contracts CNS pericytes: Role of voltage-dependent calcium channels. Proceedings of the National Academy of Sciences. 2023;120(9):e2216421120.

88. Longden TA, Hill-Eubanks DC, Nelson MT. Ion channel networks in the control of cerebral blood flow. Journal of Cerebral Blood Flow & Metabolism. 2016;36(3):492–512.

89. Dabertrand F, Krøigaard C, Bonev AD, Cognat E, Dalsgaard T, Domenga-Denier V, et al. Potassium channelopathy-like defect underlies early-stage cerebrovascular dysfunction in a genetic model of small vessel disease. Proceedings of the National Academy of Sciences. 2015;112(7):E796–E805.

90. Geary GG, Krause DN, Duckles SP. Estrogen reduces myogenic tone through a nitric oxide-dependent mechanism in rat cerebral arteries. American Journal of Physiology-Heart and Circulatory Physiology. 1998;275(1):H292–H300.

91. Skarsgard P, Van Breemen C, Laher I. Estrogen regulates myogenic tone in pressurized cerebral arteries by enhanced basal release of nitric oxide. American Journal of Physiology-Heart and Circulatory Physiology. 1997;273(5):H2248–H2256.

92. Jackson WF. Myogenic tone in peripheral resistance arteries and arterioles: the pressure is on! Frontiers in physiology. 2021;12:699517.

93. Cipolla MJ, Sweet J, Chan SL, Tavares MJ, Gokina N, Brayden JE. Increased pressure-induced tone in rat parenchymal arterioles vs. middle cerebral arteries: role of ion channels and calcium sensitivity. Journal of applied physiology. 2014;117(1):53–59.

94. Sancho M, Kyle BD. The large-conductance, calcium-activated potassium channel: a big key regulator of cell physiology. Frontiers in Physiology. 2021;12:750615.

95. Dabertrand F, Nelson MT, Brayden JE. Ryanodine receptors, calcium signaling, and regulation of vascular tone in the cerebral parenchymal microcirculation. Microcirculation. 2013;20(4):307–316.

96. Grutzendler J, Nedergaard M. Cellular control of brain capillary blood flow: in vivo imaging veritas. Trends in neurosciences. 2019;42(8):528–536.

97. Joseph A, Guevara-Torres A, Schallek J. Imaging single-cell blood flow in the smallest to largest vessels in the living retina. elife. 2019;8:e45077.

98. Klein SP, De Sloovere V, Meyfroidt G, Depreitere B. Differential hemodynamic response of pial arterioles contributes to a quadriphasic cerebral autoregulation physiology. Journal of the American Heart Association. 2022;11(1):e022943.

99. Harper SL, Bohlen HG, Rubin MJ. Arterial and microvascular contributions to cerebral cortical autoregulation in rats. American Journal of Physiology-Heart and Circulatory Physiology. 1984;246(1):H17–H24.

100. Hoffman WE, Edelman G, Kochs E, Werner C, Segil L, Albrecht RF. Cerebral autoregulation in awake versus isoflurane-anesthetized rats. Anesthesia & Analgesia. 1991;73(6):753–757.

101. Dalkara T, Irikura K, Huang Z, Panahian N, Moskowitz M. Cerebrovascular responses under controlled and monitored physiological conditions in the anesthetized mouse. Journal of Cerebral Blood Flow & Metabolism. 1995;15(4):631–638.

102. Wei HS, Kang H, Rasheed IYD, Zhou S, Lou N, Gershteyn A, et al. Erythrocytes are oxygen-sensing regulators of the cerebral microcirculation. Neuron. 2016;91(4):851–862.

103. Sancho M, Klug NR, Mughal A, Koide M, Huerta de la Cruz S, Heppner TJ, et al. Adenosine signaling activates ATP-sensitive K+ channels in endothelial cells and pericytes in CNS capillaries. Science Signaling. 2022;15(727):eabl5405.

104. Fernández-Klett F, Offenhauser N, Dirnagl U, Priller J, Lindauer U. Pericytes in capillaries are contractile in vivo, but arterioles mediate functional hyperemia in the mouse brain. Proceedings of the National Academy of Sciences. 2010;107(51):22290–22295.

105. Institoris A, Vandal M, Peringod G, Catalano C, Tran CH, Yu X, et al. Astrocytes amplify neurovascular coupling to sustained activation of neocortex in awake mice. Nature Communications. 2022;13(1):7872.

106. Grubb S, Cai C, Hald BO, Khennouf L, Murmu RP, Jensen AG, et al. Precapillary sphincters maintain perfusion in the cerebral cortex. Nature communications. 2020;11(1):1–12.

107. Cai C, Zambach SA, Grubb S, Tao L, He C, Lind BL, et al. Impaired dynamics of precapillary sphincters and pericytes at first-order capillaries predict reduced neurovascular function in the aging mouse brain. Nature aging. 2023;3(2):173–184.

108. Pries AR, Secomb T, Gessner T, Sperandio M, Gross J, Gaehtgens P. Resistance to blood flow in microvessels in vivo. Circulation research. 1994;75(5):904–915.

109. Pries AR, Secomb TW. Microvascular blood viscosity in vivo and the endothelial surface layer. American Journal of Physiology-Heart and Circulatory Physiology. 2005;289(6):H2657–H2664.

110. Pries A, Ley K, Claassen M, Gaehtgens P. Red cell distribution at microvascular bifurcations. Microvascular research. 1989;38(1):81–101.

111. Chen BR, Kozberg MG, Bouchard MB, Shaik MA, Hillman EM. A critical role for the vascular endothelium in functional neurovascular coupling in the brain. Journal of the American Heart Association. 2014;3(3):e000787.

112. Longden TA, Dabertrand F, Koide M, Gonzales AL, Tykocki NR, Brayden JE, et al. Capillary K+-sensing initiates retrograde hyperpolarization to increase local cerebral blood flow. Nature neuroscience. 2017;20(5):717–726.

113. Girouard H, Bonev AD, Hannah RM, Meredith A, Aldrich RW, Nelson MT. Astrocytic endfoot Ca2+ and BK channels determine both arteriolar dilation and constriction. Proceedings of the National Academy of Sciences. 2010;107(8):3811–3816.

114. Zhang D, Ruan J, Peng S, Li J, Hu X, Zhang Y, et al. Synaptic-like transmission between neural axons and arteriolar smooth muscle cells drives cerebral neurovascular coupling. Nature Neuroscience. 2024; p. 1–17.

115. Klein S, De Sloovere V, Meyfroidt G, Depreitere B. Autoregulation assessment by direct visualisation of pial arterial blood flow in the piglet brain. Scientific reports. 2019;9(1):13333.

116. Takahashi T, Nagaoka T, Yanagida H, Saitoh T, Kamiya A, Hein T, et al. A mathematical model for the distribution of hemodynamic parameters in the human retinal microvascular network. Journal of biorheology. 2009;23:77–86.

117. Bertolini M, Causin P, Turrini C. A mathematical characterization of anatomically consistent blood capillary networks. Journal of Mathematics in Industry. 2023;13(1):1–17.

